# Age-associated Senescent - T Cell Signaling Promotes Type 3 Immunity that Inhibits Regenerative Response

**DOI:** 10.1101/2021.08.17.456641

**Authors:** Jin Han, Christopher Cherry, Joscelyn C. Mejias, Anna Ruta, David R. Maestas, Alexis N. Peña, Helen Hieu Nguyen, Brenda Yang, Elise Gray-Gaillard, Natalie Rutkowski, Kavita Krishnan, Ada J. Tam, Elana J. Fertig, Franck Housseau, Sudipto Ganguly, Erika M. Moore, Drew M. Pardoll, Jennifer H. Elisseeff

**Affiliations:** Translational Tissue Engineering Center, Wilmer Eye Institute and Department of Biomedical Engineering, Johns Hopkins University; Baltimore, MD, USA; Bloomberg∼Kimmel Institute for Cancer Immunotherapy, Sidney Kimmel Comprehensive Cancer Center, Johns Hopkins University School of Medicine; Baltimore, MD, USA; Department of Biomedical Engineering and Institute for Cell Engineering, Johns Hopkins University School of Medicine; Baltimore, MD, USA; Department of Oncology, Johns Hopkins University School of Medicine; Baltimore, MD, USA; Department of Applied Mathematics and Statistics, Johns Hopkins University; Baltimore, MD, USA; Department of Materials Science and Engineering, University of Florida; Gainesville, FL USA

## Abstract

Aging is associated with immunological changes that compromise response to infections and vaccines, exacerbate inflammatory diseases and could potentially mitigate tissue repair. Even so, age-related changes to the immune response to tissue damage and regenerative medicine therapies remain unknown. Here, we characterized how aging induces senescence and immunological changes that inhibit tissue repair and therapeutic response to a clinical regenerative biological scaffold derived from extracellular matrix. Tissue signatures of inflammation and interleukin (IL)-17 signaling increased with injury and treatment in aged animals, and computational analysis uncovered age-associated senescent-T cell communication that promotes type 3 immunity in T cells. Local inhibition of type 3 immune activation using IL17-neutralizing antibodies improved healing and restored therapeutic response to the regenerative biomaterial, promoting muscle repair in older animals. These results provide insights into tissue immune dysregulation that occurs with aging that can be targeted to rejuvenate repair.

---

Aging is associated with decreased tissue function and a compromised response to tissue damage that leads to longer recovery and frequently dysfunctional tissue repair regardless of tissue type^1-3^. Reduced healing capacity with increasing age was recognized as early as 1916^4^. Consistent with the variability in biological signatures of aging, the variability in time required for tissue repair and the variability in resulting tissue quality increases with age in both preclinical models and patients^5-7^. Multi-omic analyses of cellular and molecular profiles of organisms over lifespan implicate changes in gene expression, metabolism, DNA methylation and epigenetic factors in age-associated pathologies including impaired wound healing^2,3^. Recovering tissue repair capacity that is lost with aging represents a significant medical challenge. While murine models of tissue repair and regeneration have brought forward transformational learnings, the vast majority of studies have been done in young mice – the equivalent of human adolescents. However, most ailments clinically affecting tissue health in humans are in the elderly demographic.

Regenerative medicine and tissue engineering approaches are designed to enhance repair and restore tissue function. While many patients needing regenerative medicine technologies are older, the influence of age-related physiological changes on regenerative medicine therapeutic responses remains unexplored. In fact, age-related changes may be, in part, related to the disappointing clinical translation and efficacy of tissue engineering technologies and should be considered in their design. The composition and phenotype of cells responding to tissue damage change with age. In skin wounds, the number of fibroblasts responding to injury is greater in older mice and the fibroblasts have reduced phenotypic heterogeneity compared to the wounds in younger counterparts^8^. In the case of muscle tissue, the number and activity of muscle stem cells decrease with age leading to sarcopenia and impaired muscle healing after injury^9^. However, the functionality of aged muscle stem cells can be restored *ex vivo* to recover healing capacity after re-injection *in vivo*, suggesting that endogenous repair capacity is retained but the aging tissue environment impedes repair^9^. Similarly, repair in the aging retina could be restored by targeting age-related epigenetic changes^10^, again suggesting that regeneration capacity remains with increasing age despite decreased cell numbers and inhibitory factors.

Classical regenerative medicine strategies utilize stem cells, growth factors and biomaterials alone or in combination to promote tissue development^11^. More recently, the role of the immune system in tissue repair is being recognized as a key factor in determining healing outcomes leading to the introduction of immunomodulation as a new therapeutic modality in regenerative medicine technology design. However, there are also numerous age-related changes that occur in the immune system that may impede a regenerative therapeutic response^12^. Age-related immune changes have been primarily studied in the context of infectious disease, chronic inflammatory conditions, vaccine efficacy and more recently cancer immunotherapy efficacy, but they may also negatively impact the response to tissue damage and regenerative medicine therapies^13^. For example, T cell numbers decrease with aging and there is a myeloid shift in the bone marrow^14,15^. Furthermore, there are composition changes in the T cell compartment with aging that include increased CD8^+^ T cells, reduced naïve CD4 ^+^ T cells, and increased effector CD4^+^ T cells, which altogether may compromise a pro-regenerative immune response and tissue repair^14^.

Here, we investigated how immunological changes associated with aging impact the response to muscle injury and limit the regenerative capacity of a therapeutic biological scaffold. We show that age-related increases in hardwired chronic interleukin (IL)-17 production by innate and adaptive elements of the immune system combined with dysfunctional stromal interactions antagonize tissue regeneration while supporting fibrosis and adipogenesis. IL17 blockade thus therapeutically reverses this age-associated impairment. Targeting age-associated immunological changes that inhibit a regenerative response may enable recovery of a therapeutic response and restoration of tissue repair capacity in older organisms.

## Aging reduces type 2 immune and tissue repair responses to regenerative biomaterials

To evaluate the impact of aging on the immune response and resulting repair efficacy, we characterized the response to an extracellular matrix (ECM) biomaterial in a muscle wound in young (6 week) and old (72 week) mice (**Fig. 1a**). ECM biomaterials, derived from the matrix of different porcine and human tissue types^16^, are used for tissue repair in multiple clinical indications^17,18^ and are more easily delivered than cells and growth factors^19-21^. In this study, we utilized clinically available^22^ ECM from porcine small intestinal submucosa (SIS) as an example biological material to model age-related differences in response. Previous research showed that muscle repair requires a type 2 immune response with IL4 signaling^23,24^. Application of an ECM biomaterial in a muscle injury increased IL4 expression and promoted repair in part by increasing recruitment of IL4 producing eosinophils and type 2 helper T (Th2) cells^25^.

**Figure 1.**
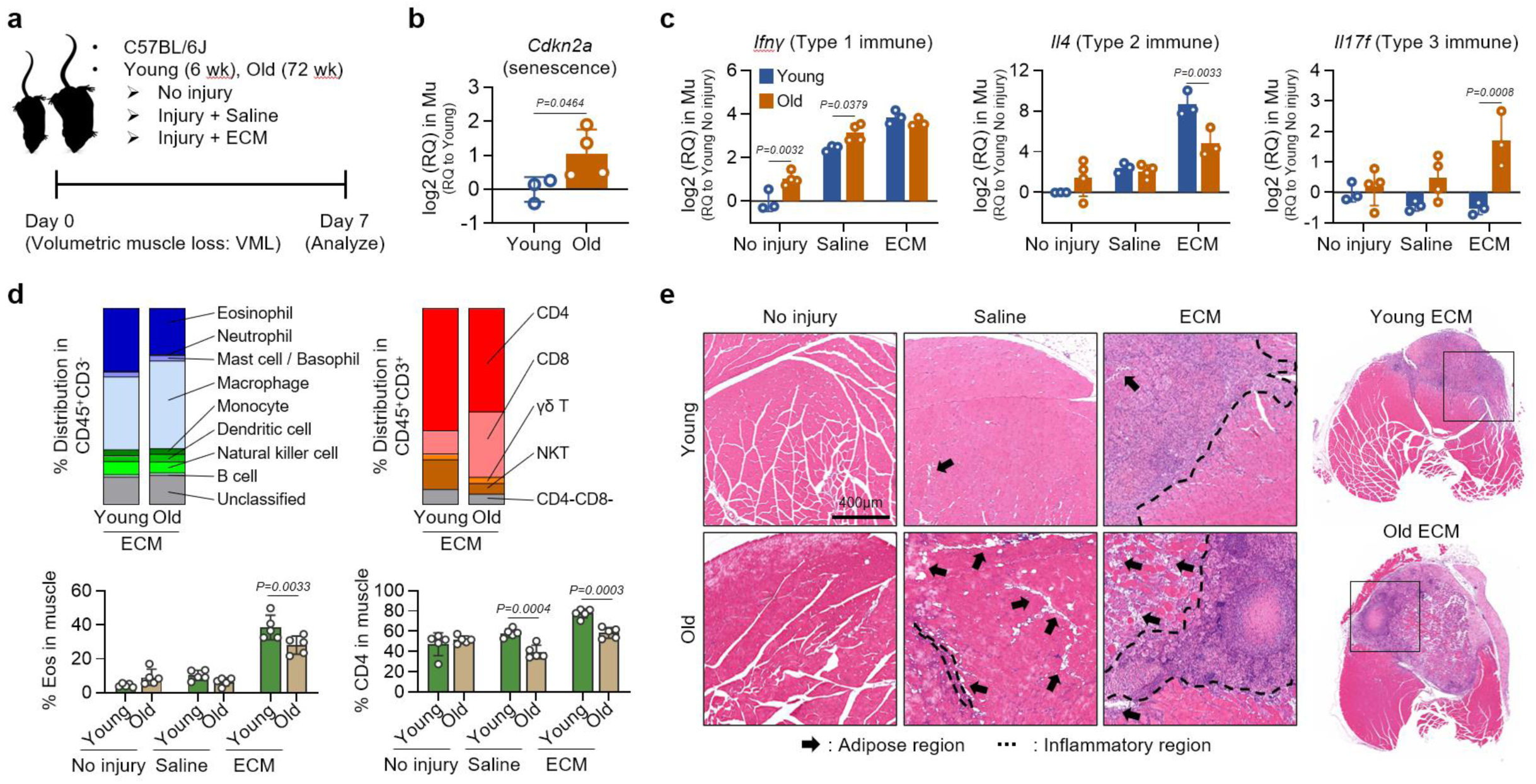
Aging alters immune and stromal response to regenerative ECM biomaterials in muscle. **a**, Schematic illustration of experimental design including no injury control, volumetric muscle loss injury (VML) treated with saline and VML treated with ECM in young (6 wk) and old (72 wk). **b**, Quantification of a senescence gene *Cdkn2a* (*p16*) in muscle without injury or ECM. **c**, Quantification of type 1, type 2 or type 3 immune response-related genes in muscle 1 week after injury or ECM treatment. **d**, Quantification of immune cells in muscle 1 week after treatments as determined by spectral flow cytometry (top) and quantification of eosinophils and CD4 T cells (bottom), a potential source of type 2 immune response. Indicated cell population represents an average value of n=5 per group. Eosinophils are presented as % of CD11b^+^, and CD4 as % of CD3^+^γδ^-^NK^-^. **e**, Transverse section of the quadricep muscle 1 week after injury or ECM stained with H&E. The black arrow indicates the ectopic adipogenesis region, and the dotted line demonstrates immune cell infiltrated area. Unpaired two-tailed t-test (**b**), two-way ANOVA with Sidak’s multiple comparisons test within the treatment group (**c**-**d**). For all bar graphs, data are mean ± s.d.

Aged muscle had a higher expression of a senescent-associated gene *Cdkn2a*, the gene that encodes p16^INKA^, even before injury (**Fig. 1b**), and significantly altered the cytokine gene expression profiles 1 week after volumetric muscle loss (VML) injury or ECM implantation (**Fig. 1c**). Expression of type 1 immune response by *Ifnγ* in muscle tissue was significantly higher in old animals than young animals after injury. Instead, type 2 immune response, represented by *Il4* gene expression, increased after injury with further significant increases after ECM treatment in young animals. However, *Il4* expression did not increase after injury in old mice and its level of expression after ECM treatment was significantly lower than the young animals. Instead, ECM implantation in old mice increased the gene expression of type 3 immune response, *Il17f* (*Il17a* gene expression was not detected in the muscle tissue of young or aged mice). Other inflammatory genes, including *Il23a, Il6, Il1b*, and *S100a4*, all increased similarly in both young and old mice 1 week after injury or ECM implantation (**Fig. S1**).

Next, using multiparametric spectral flow cytometry (**Fig. S2**), we found injury and ECM treatment resulted in distinct changes in immune and stromal cell responses in an aging muscle environment (**Fig. 1d** and **Fig. S3-5**). A robust cellular response to injury and ECM implant occurred in both young and aged animals, but there were significant differences in the composition of the immune and stromal compartments. Eosinophils, a major source of regenerative type 2 response, increased in numbers 1 week after ECM treatment in both young and aged animals, however, their general percentage was significantly lower in old mice compared to the younger counterparts (**Fig. 1d**). Additionally, the adaptive T cell immune response to muscle injury and ECM treatment changed with age as more CD8 cells responding to injury and ECM in the aged animals compared to the CD4 cell response in the young (**Fig. 1d** and **Fig. S5**). These data suggested that ECM-induced recruitment and expansion of cell sources for type 2 immune responses, such as eosinophils and CD4 T cells, was significantly limited in aging tissue environment. Interestingly, the CD45^-^ population, which included fibroblasts, endothelial and other stromal cells, notably increased in numbers 1 week after ECM implantation in aged animals compared to young animals (*p*=0.0057; **Fig. S4**).

The differences in immune cell recruitment and cytokine expression in muscle correlated with changes in tissue repair observed histologically (**Fig. 1e** and **Fig. S6**). One week after injury or ECM treatment, there was a significant cell infiltration in both young and aged mice, with ECM treatment further increasing cellular infiltration and collagen deposition as visualized by Masson’s Trichrome staining. There was also excessive adipose tissue in the muscle of aged animals compared to the younger counterparts, which increased further with ECM treatment (**Fig. 1e**). Together, these data demonstrated that aging limited the regenerative immune response to ECM material, and instead promoted inflammatory type 3 immunity, fibrosis, and adipogenesis.

### Single cell analysis reveals age-specific immune and stromal response after injury and regenerative medicine treatment

To further identify aging signatures of injury and therapeutic response to a regenerative biomaterial therapy, we performed single cell RNA sequencing (scRNA-seq) on CD45^+^-enriched cells isolated from the muscle tissue 1 week after injury or ECM implantation (**Fig. 1a**). We found multiple distinct cell clusters in the merged samples (**Fig. 2a** and **Fig. S7-9**) that included myeloid cells, T cells, granulocytes, fibroblasts, endothelial/pericytes, and skeletal muscle cells. We observed two different macrophage subtypes containing Mrc1^hi^ macrophage (Mrc1^hi^ Mc; *Mrc1*^hi^*Ccl8*^hi^) and Arg1^hi^ macrophage (Arg1^hi^ Mc; *Arg1*^hi^*Mmp12*^hi^) and detected multiple clusters in the fibroblast population consisting of generic fibroblast (Gen Fib; *Col1a1*^hi^*Col3a1*^hi^), Mgp^hi^ fibroblast (Mgp^hi^ Fib; *Mgp*^hi^*Apod*^hi^), and Pi16^hi^ fibroblast progenitor (Pi16^hi^ Fib; *Pi16*^hi^*Cd34*^hi^). Additionally, we identified generic myeloid cell (Gen Myl; *Adgre1*^lo^*Ccr2*^hi^*Cd74*^hi^), granulocyte cluster 1 (Gr-1; *S100a8*^hi^*Il1f9*^hi^), granulocyte cluster 2 (Gr-2; *Ngp*^hi^*Camp*^hi^), T cells (*Trbc2*^hi^), CD209 dendritic cell (CD209 DC; *Cd209a*^hi^), a combination of endothelial cells and pericytes (Endo/Peri; *Fabp*^hi^*Rgs*5^hi^), and skeletal muscle cells (SMC; *Acta1*^hi^*Myh4*^hi^).

**Figure 2.**
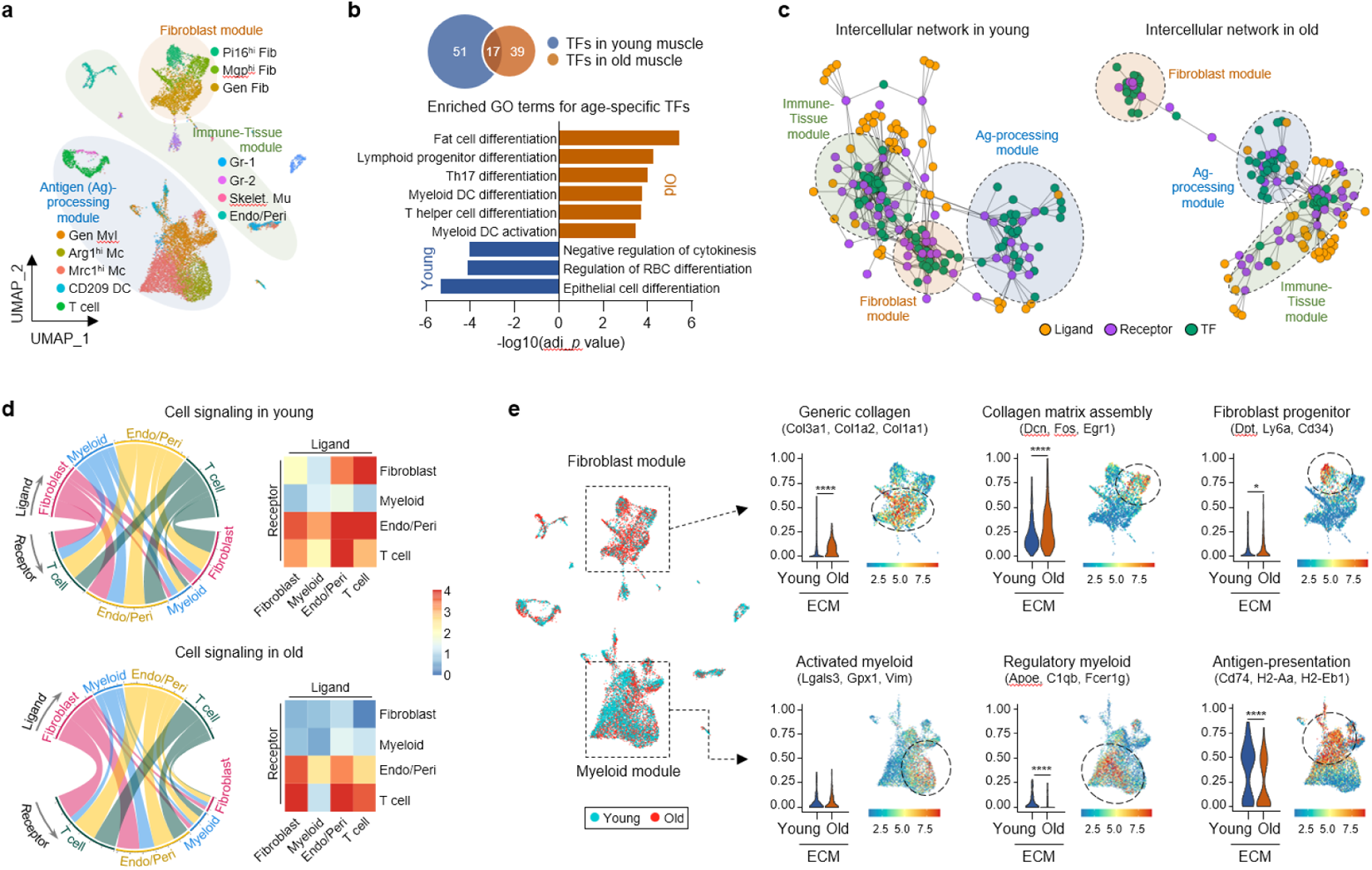
Immune-stromal communication in muscle is impaired with aging. **a**, UMAP overview of cell clusters identified using scRNA-seq dataset on muscle from young and old animals 1 week after injury or treatment (3 mice pooled for each condition). Clusters were self-assembled into three signaling modules enriched in fibroblast, antigen-processing, and immune-tissue clusters. **b**, Gene set enrichment analysis of age-specific transcription factors (TF) in muscle. Adjusted *p* values (log 10) of significant GO terms are shown. **c**, Age-specific global signaling network in muscle. Three distinct modules are labelled on the basis of enrichment of receptors and transcription factors by each cluster. **d**, Chord plot (left) and heatmap (right) of predicted cluster-cluster signaling for selected clusters determined by Domino for young or old muscle. Pairwise interactions are shown between ligand and receptor genes expressed by fibroblast, myeloid/macrophage, endothelial/pericyte, and T cell clusters. The values shown are the summed z-scored expression values for ligands (in the ligand-cluster) targeting receptors predicted to be activated in the receptor-cluster. Higher values indicate increased expression of ligands predicted to be active for a given receptor cluster. The width of the chord shows the strength of the interaction. **e**, NMF-CoGAPS analysis of fibroblast populations (top) and myeloid/macrophage populations (bottom) in muscle 1 week after ECM treatment. Region of cells expressing high levels of the gene sets are circled. *p* values in NMF-CoGAPS were determined using Mann-Whitney U test and adjusted with false discovery rate correction for multiple testing. \**p*<0.05, and \*\*\*\**p*<0.0001.

We then observed age-dependent changes in cell clusters from ECM-treated muscle injuries in young and aged animals (**Fig. S10**). Granulocyte cluster Gr-2 enriched with *Camp* expression, a gene associated with early transcriptional states of neutrophils^26,27^ that induce migration of eosinophils^28,29^, increased after injury and ECM implantation only in young animals, supporting the increased eosinophil migration in young muscle found by flow cytometry (**Fig. 1d**). The skeletal muscle cell cluster also increased with ECM treatment only in young animals, further suggesting that ECM increased myogenic activity primarily in young mice. On the other hand, the CD209 dendritic cell cluster, whose cytokines are known to promote Th17 phenotypes in T cells^30-33^, increased with ECM only in old mice.

### Aged animals exhibit impaired immune-stromal communication required for tissue repair

Tissue repair requires removal of debris, mobilization of stem cells, vascularization, and secretion and organization of tissue-specific extracellular matrix that is coordinated through complex immune-stromal cell interactions. To further probe the differences in the young and aged tissue environment after trauma and biomaterial application, we applied Domino to model cell-cell communication patterns using the data obtained from scRNA-seq. Domino is a computational tool that identifies condition-specific intercellular signaling dynamics based on transcription factor (TF) activation, which is surmised based on regulon expression with SCENIC gene regulatory network analysis^34^, along with receptor (R) and ligand (L) expression independent of clusters^35^. Domino constructs a signaling network connecting TF-R-L, which are specifically predicted to be active in the dataset. TF-R connections are determined by examining correlation between R expression and TF activation scores across all cells in the dataset, identifying TF-R pairings with grouped increases of expression and activation in target cell populations. R-L pairs are then determined for target receptors through the CellphoneDB2 database.

To understand the biological significance of the age-specific TFs, we first performed gene set enrichment analysis on the enriched gene ontology (GO) terms to classify the biological processes in which they function (**Fig. 2b**). The young-specific TFs were associated with regulation of red blood cell differentiation or cytokinesis, and epithelial cell differentiation. On the other hand, old animal-specific TFs were enriched for fat cell differentiation, Th17 differentiation and myeloid dendritic cell differentiation pathways. Signaling associated with Th17 differentiation is known to negatively regulate eosinophil recruitment and IL4 expression^36-38^, suggesting that age-associated immunological skewing in the TFs may be responsible for the impaired Th2 response to the regenerative ECM biomaterial in old animals. Additionally, enriched genes related to fat cell and myeloid dendritic cell differentiation in old animals correlated with our histological (**Fig. 1e**) and scRNA-seq (**Fig. S10**) data.

Next, we observed cell-cell communication patterns as predicted by Domino. In both young and old animals, a force directed diagram of the TF-R-L signaling network self-assembled into three signaling modules enriched in fibroblast, antigen processing and immune-tissue clusters (**Fig. 2c**). Each module indicates signaling pathways with similarly enriched activation in specific cell types. Increased module density and decreased connection across modules both indicate a group of highly correlated signaling patterns expressed in a specific cell population (both receptors and transcription factors). A complete list of the TFs, and the receptors corresponding to the activated TFs is provided in **Fig. S11-13**. Increased connectivity between the fibroblast and immune-tissue modules in the signaling network of young muscle indicated some level of immunological properties in fibroblasts and their plasticity in signaling (**Fig. 2c**). On the other hand, the fibroblasts in the aged tissue appeared to lose immunological properties and the narrow localization of the activated TFs suggested reduced heterogeneity. Single cell cluster ligand - receptor expression and correlation identified in further detail age-associated changes in inter-cluster signaling (**Fig. 2d** and **Fig. S14**). We observed that aging disrupted the ligand-to-receptor signaling between T cell and Endo/Peri, and T cell and fibroblast, both of which were active in young mice. Myeloid communication with T cells also diminished with aging. Overall, the communication predicted between the receptors on fibroblasts and the ligands expressed in all other clusters decreased in old mice.

To better elucidate the age-related gene signatures in fibroblast and myeloid clusters, we then utilized a Bayesian non-negative matrix factorization (NMF) algorithm termed coordinate gene activity in pattern sets (CoGAPS) to capture additional gene sets representing cellular processes from the single cell dataset independent of changes in cell clusters^39^ (**Fig. 2e**). NMF is an alternative method to infer expression patterns that can span multiple clusters, reflective of biological processes^40^, with the Bayesian framework of CoGAPS having additional sparsity constraints ideal for scRNA-seq analysis. In the present dataset, CoGAPS found gene signatures of collagen matrix assembly dominant in the aged muscle (**Fig. 2e, upper panel** and **Fig. S15**), supporting the increased fibrosis observed histologically (**Fig. S6**). Specifically, a set of genes related to collagen assembly, such as *Dcn, Fos*, and *Egr1*^41-43^, was more prominent in old fibroblasts with ECM treatment showing the highest levels. Similarly, genes associated with generic collagen, such as *Col3a1* and *Col1a1/2*, or those with fibroblast progenitors, represented by *Dpt, Ly6a*, and *Cd34*, were all more dominantly expressed in fibroblasts from old animals.

CoGAPS also highlighted the gene profiles in the myeloid and macrophage cells that were more dominant in young animals (**Fig. 2e, lower panel** and **Fig. S16**). Genes associated with activated (*Lgals3, Gpx1, Vim*)^44,45^ or regulatory (*Apoe, C1qb, Fcer1g*)^46,47^ myeloid cells were highly expressed in young muscle compared to aged muscle. While the treatment with ECM increased the expression of MHCII-associated genes (*Cd74, H2-Aa, H2-Eb1*) in both young and old animals, their gene signatures were much more distinct within the young. This suggests that even though some cell clusters increased with ECM implantation in a similar manner between young and old mice, they may have different gene signatures that impact functional outcomes.

Altogether, the flow cytometry and single cell analysis demonstrate that key immune populations involved in muscle repair and a regenerative therapeutic response, such as eosinophils and CD4 T cells, decrease with aging. Furthermore, aging promotes pro-inflammatory cells such as CD8 T cells, increases type 3 immune responses, and develops unique fibrosis signatures in response to regenerative treatments while at the same time decreasing immune activities in myeloid clusters relevant for tissue repair *via* antigen presentation and immune cell mobilization.

### Aging induces a systemic type 3 adaptive immune response to injury and biomaterial therapy

To further evaluate age-associated adaptive immunity, we then analyzed the baseline proximal inguinal lymph node (iLN). We first compared the gene expression profile of iLNs from naïve (no injury) young and old mice using Nanostring (**Fig. 3a** and **Fig. S17-18**) and flow cytometry (**Fig. 3b** and **Fig. S19**). Aged lymph nodes expressed higher expression of NF-κb/TNFα-, Fc receptor- or Th17-associated gene sets, all of which are potent inducers of Th17-mediated inflammation^48^ (**Fig. 3a** and **Fig. S17**). Pathway scoring also suggested that aging promoted T cell activation in Th17 differentiation or Th17-biology related gene expression (**Fig. S18**). Flow cytometric analysis further supported a type 3-skewed adaptive immune environment in the aging lymph node with a significantly higher proportion of IL17A-producing CD4 or γδ T cells compared to the young (**Fig. 3b** and **Fig. S19**). While the percentage of Th1 (IFNγ^+^ CD4) and Th17 (IL17A^+^ CD4) cells both increased with aging, the γδ T cells switched phenotype from type 1 (IFNγ^+^) to type 3 (IL17A^+^) immunity.

**Figure 3.**
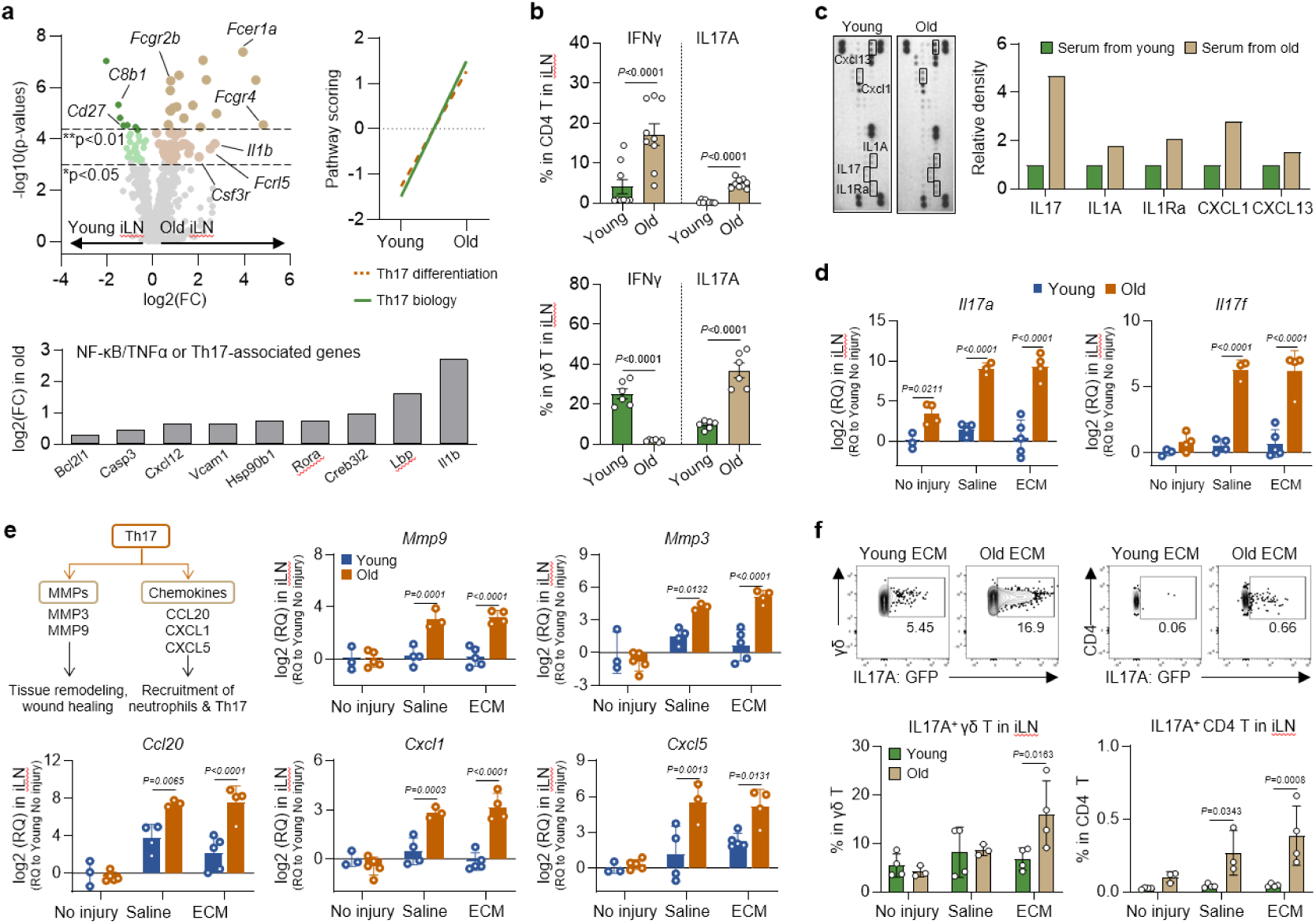
Aging induces a Th17-associated immune skewing, and injury and ECM treatment promote local and systemic type 3 immune response that inhibits tissue repair. **a**, Volcano plot of genes expressed in aged lymph node normalized to those in young lymph node (top left). Gene pathway scoring for type 17 helper T cell (Th17) pathways (top right) or differentially expressed genes for NF-κb/TNFα or Th17-associated pathways (bottom) based on Nanostring analysis are shown. **b**, Multiparametric flow cytometry quantification of IFNγ^+^ or IL17A^+^ CD4 or γδ T cells in the lymph nodes from young or aged animals without injury or treatment. **c**, Representative images (left) and quantitative analysis (right) of the proteome profiler performed on serum from young and old mice without the treatments (3 mice pooled for each condition). Protein molecules with significant differences in pixel densities compared to young animals are labeled and quantified using imageJ. **d**, Quantification of Th17 genes in lymph node 1 week after injury or ECM treatment. **e**, Quantification of Th17-associated genes in the lymph nodes. A schematic of the downstream production of matrix metalloproteinase (MMP) and chemokine from Th17 is shown (top left). **f**, Representative images (top) and quantification (bottom) of flow cytometry data comparing IL17A^+^ γδ^+^ or CD4^+^ T cells between young and old animals in the lymph node 1 week after injury or ECM treatment. Unpaired two-tailed t-test (**b**), two-way ANOVA with Sidak’s multiple comparisons test within the treatment group (**d**-**f**). For all bar graphs, data are mean ± s.e.m (**b**) or s.d. (**d**-**f**).

To further assess the systemic immune changes with aging, we analyzed serum proteins using proteome profiler between young and old mice (**Fig. 3c** and **Fig. S20**). In addition to increased IL17 in the serum of aged mice, there was also increased level of the B cell chemoattractant factor CXCL13, which also increased after injury only in old mice. Additionally, IL1A, a potent inducer of IL17 from T cells^49^, and CXCL1, one of IL17-induced chemokines, also increased in serum from the aged mice. IL23, a protein that can expand Th17 cells^50^ increased in response to injury only in old mice.

We then evaluated the regional response to injury and ECM treatment in the lymph nodes from young and aged mice (**Fig. 3d-f** and **Fig. S21**). First, the gene expression for helper T cell cytokines in iLNs correlated with the immune profiles found in the muscle tissue (**Fig. 3d**). Strikingly, genes associated with type 3 immune response, *Il17a* and *Il17f*, significantly increased only in aged animals after injury or ECM treatment. Thereafter, we further analyzed the expression of the downstream genes related to Th17; *Mmp3, Mmp9, Cxcl1, Ccl20*, and *Cxcl5*^51^. Similar to *Il17a* and *Il17f*, the expression of the downstream genes occurred at low levels in naïve young and aged animals, but significantly increased only after injury or ECM treatment in old mice (**Fig. 3e**), suggesting that injury further triggered a type 3 immune response beyond the baseline in aging muscle tissue. Multiparametric flow cytometry analysis of iLNs of IL17A-IRES-GFP-KI (IL17A-GFP) mice further confirmed this trend with increase in percentages of IL17A^+^ γδ or CD4 T cells after ECM treatment in aged mice (**Fig. 3f**). Collectively, these data show that injury and ECM implantation in aged animals trigger a systemic type 3 adaptive immune response that may be responsible for the decreased regenerative response with aging.

### Transfer learning identifies changes in senescent cell - T cell communication with aging

To identify the signaling components that maybe responsible for impaired wound healing in old animals, and evaluate the mechanistic of type 3 immune skewing with aging, we investigated age-associated TF activation in T cells and their communication in the signaling networks. We developed a transfer learning algorithm to identify senescent cells in single cell datasets using a senescence signature derived from a p16-Cre;Ai14 reporter model^52^. Transfer learning identified 3 stromal clusters expressing a high senescent signature, with the Mgp^hi^ fibroblast cluster showing the highest senescence signature enrichment (**Fig. 4a** and **Fig. S22**). While transfer learning identified the Mgp^hi^ fibroblast cluster as being senescent in both the young and old animals, the senescence-associated secretory phenotype (SASP) differed and the SASP in old animals was enriched with *Tgfβ3, Col11a1*, and *Timp1*. The SASP of the senescent cluster in young and old animals is provided in **Fig. S22**.

**Figure 4.**
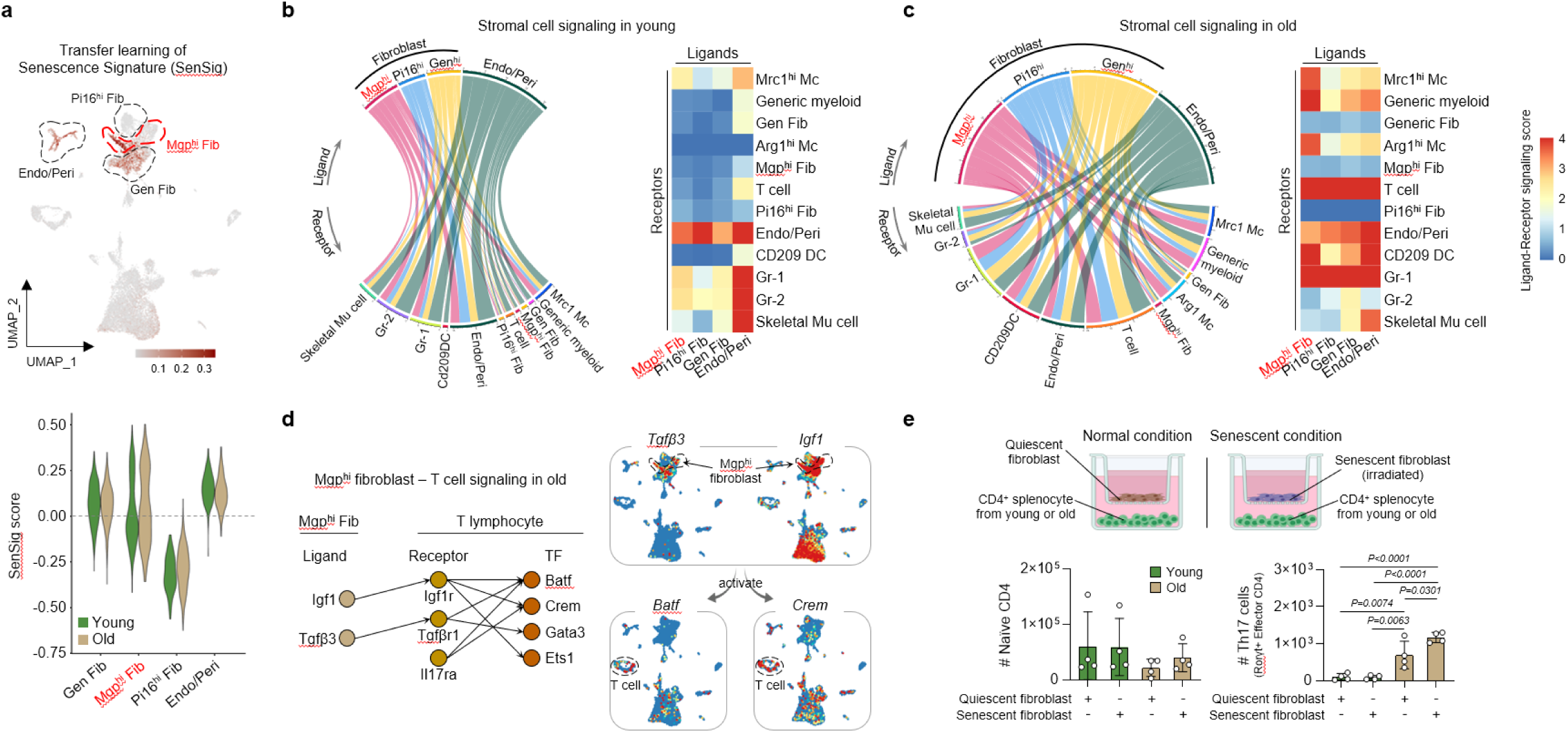
Age-associated signaling communication between senescent stromal cells and immune cells induces type 3 immune skewing in old animals. **a**, Senescence scores for cells shown on UMAP (top) and violin plot grouped by cluster and age (bottom). CD45^-^ stromal clusters are marked with dotted line, and a cluster with the highest senescence signature is marked red. (**b**-**c**) Chord plot (left) and heatmap (right) of predicted cluster-cluster signaling for selected clusters in young (**b**) or old (**c**) muscle. Pairwise interactions are shown between ligands of CD45^-^ clusters, fibroblasts and endothelial/pericyte, and receptors of all other clusters. The values shown are the summed z-scored expression values for ligands (in the ligand-cluster) targeting receptors predicted to be activated in the receptor-cluster. Higher values indicate increased expression of ligands predicted to be active for a given receptor cluster. The width of the chord shows the strength of the interaction. **d**, Identification of ligand-receptor-transcription factor (TF) signaling network between senescent fibroblast (Mgp^hi^ fibroblast) and T cells in old animals (left), and its UMAP representation (right). SCENIC is used to estimate TF modules and activation scores. Receptor expressions are correlated with TF activation scores with exclusion of receptors present in the TF modules. Public receptor–ligand databases are used to identify ligands activating receptors. **e**, Illustration of coculture platform designed to study senescent fibroblast-T cell communication *in vitro* (top) and multiparametric flow quantification of CD4 T cells after coculture (bottom). Two-way ANOVA with Tukey’s multiple comparisons test in (**e**). For all bar graphs, data are mean ± s.d.

Next, we used Domino to identify intercellular signaling communication between the senescent fibroblast and other cell clusters (**Fig. 4b-c**). We observed the signaling coming from the senescent fibroblast (Mgp^hi^ fibroblast) was limited in young mice with minimal communication predicted between the senescent fibroblast and T cells (**Fig. 4b**). In aged mice, however, there was a large increase in signaling coming from the senescent fibroblast cluster (SASP ligand expression), which correlated with receptor expression and TF activation in innate immune cells and T cell populations (**Fig. 4c**).

To further probe the detailed signaling communication between the senescent fibroblast and T cells in old mice, and identify corresponding TF activation in the adaptive immune response, we constructed TF-R-L signaling between the two clusters (**Fig. 4d**). Domino demonstrated that the *Igf1* and *Tgfβ3* secreted from the senescent fibroblast in aging muscle target *Igf1r* or *Tgfβr1* of the T cells, and subsequently activate the TFs such as *Batf* and *Crem*, both of which have been implicated in Th17 differentiation and are also activated by the receptor *Il17ra. Batf* is one of the activator protein-1 (AP-1) proteins that controls Th17 differentiation, and is the key activator in T cell receptor activation necessary for Th17 lineage specification^53^. *Crem* is also one of the key TFs that alters IL17A production when regulated^53^, and is known to enhance RORγt accumulation on the *Il17a* promoter^54^. Together, these findings suggest the SASP signaling from the senescent fibroblast in aged muscle develops type 3 immunity in T cell response, providing a biological mechanistic of immune skewing with aging that inhibits the regenerative response to ECM materials.

To experimentally validate the computationally-predicted interactions between the senescent fibroblast and T cells with aging, we investigated whether CD4 T cells from old animals can better differentiate into Th17 cells in coculture with the senescent (irradiated) cells (**Fig. 4e**). After 5 days of coculture with the senescent fibroblasts, only the CD4 cells from old mice differentiated into Th17 cells and significantly expanded in number. There was no increase in Th17 cells when CD4 cells were isolated from young mice, suggesting that only the aged T cells are primed for Th17 differentiation in communication with the senescent cells.

### Aging CD4 T cells demonstrate a unique Th17 immune phenotype

To further evaluate the age-associated type 3 adaptive immune signatures, we evaluated whether aging CD4 T cells have a higher propensity for Th17 lineage compared to young CD4 T cells (**Fig. 5**). As expected, CD4 cells isolated from the lymph nodes and spleen of aged animals exhibited fewer naïve (CD44 ^-^CD62L^-^) but significantly higher percentages of effector (CD44^+^CD62L^-^) phenotype compared to the young (**Fig. 5a**). These effector T cells had notably higher percentages of RORγt expression. To determine if the increased numbers of Th17 cells in aged tissues was due to an increased propensity for differentiation, we isolated naïve CD4 ^+^ splenocytes from young and old animals, and cultured them in Th17 skewing conditions *in vitro* (**Fig. 5b**). Interestingly, naïve CD4 T cells from aged animals demonstrated better differentiation into effector cells and RORγt^+^ Th17 cells under skewing conditions compared to those from young animals (**Fig. 5b, right panel**). In addition to increased skewing into Th17 cells, T cells from aged animals had a unique secretory phenotype after skewing. Proteome analysis on the Th17 cells differentiated from naïve CD4 cells from aged mice showed upregulation in inflammatory cytokines, including IL12p40, a subunit for IL-23 that is required for Th17 differentiation, IL-6 family leukemia inhibitory factor (LIF), CCL5, CCL6, CCL22, VEGF and many others (**Fig. 5c**). These data suggest that naïve CD4 T cells from the aging environment differentiate into a unique Th17 phenotype, and aging Th17 cells demonstrate a different secretory profile compared to young Th17 cells with more VEGF and CCL5 secretion, both of which could play a direct role in angiogenesis during tissue regeneration.

**Figure 5.**
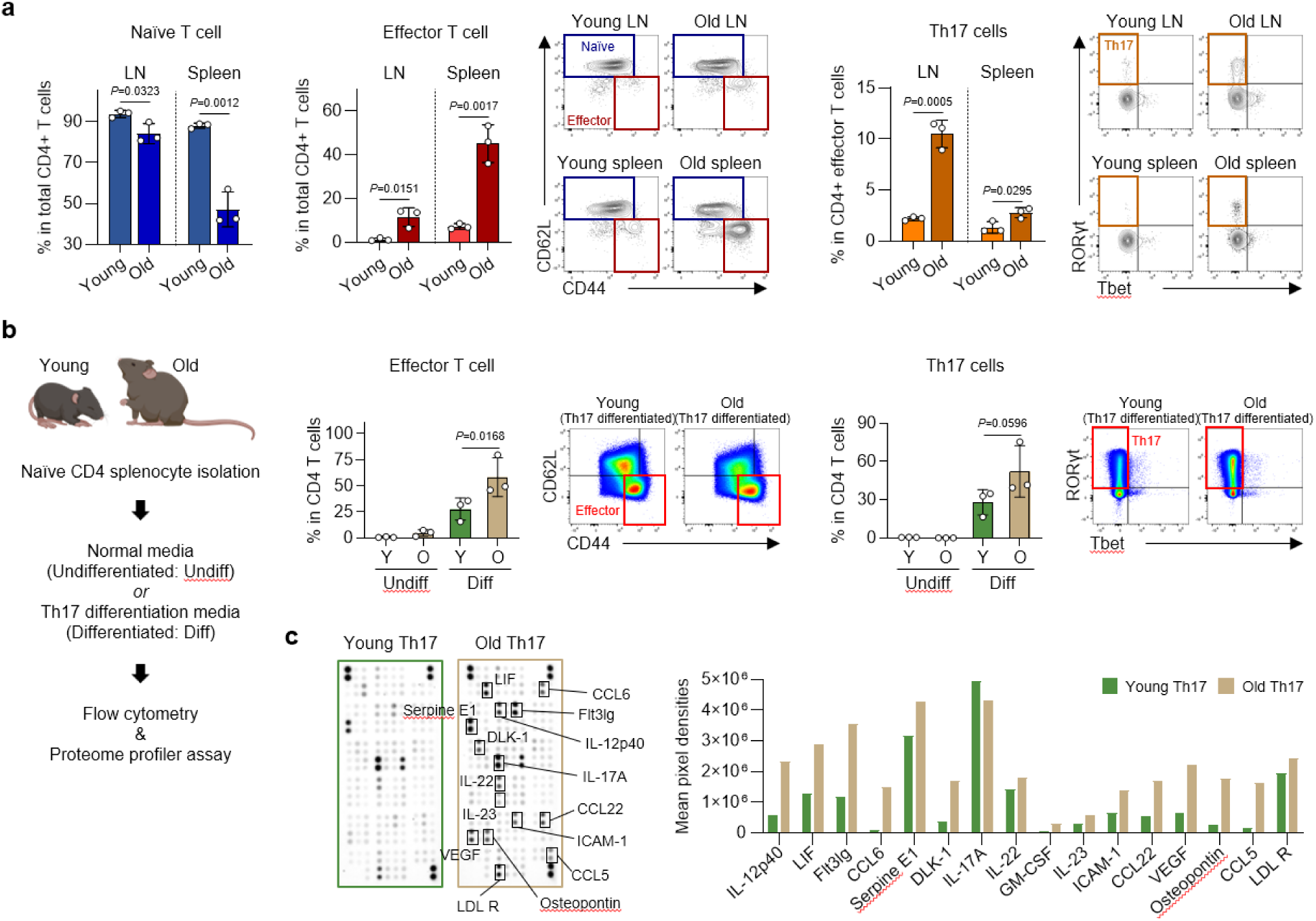
Aging is associated with increased Th17 effector T cells with a unique secretome profile. **a**, Quantification and representative plots of flow cytometry analysis on CD4 T cells isolated from lymph nodes or spleens of young and old animals. Naïve phenotype (CD4 ^+^CD44^-^CD62L^-^), effector phenotype (CD4^+^CD44^+^CD62L^-^) and Th17 cells (CD4^+^CD44^+^CD62L^-^RORγt^+^) are shown. **b**, Schematic illustration of naïve T cell isolation and Th17 differentiation *in vitro* (left) and quantification of flow cytometry analysis on the undifferentiated and differentiated CD4 T cells (right). **c**, Representative images (left) and quantification (right) of the proteome profiler performed on cell culture supernatant from Th17-differentiated CD4 T cells from young and old mice (3 samples pooled for each condition). Protein molecules with significant differences in pixel densities compared to young animals are labeled and quantified using iBright Analysis Software. Unpaired two-tailed t-test (**a**), two-way ANOVA with Sidak’s multiple comparisons test (**b**). For all bar graphs, data are s.d.

### Inhibition of IL17 rejuvenates type 2 immune response after muscle injury in aged animals

Since type 3 immune response and IL17 are associated with fibrosis^55,56^ and negatively regulate IL4 that is needed for tissue repair, we investigated whether IL17 neutralizing antibodies (αIL17) could restore IL4 expression and tissue repair that is lost with aging. We first evaluated the dosing and timing for delivery (**Fig. S23**). A minimum of three injections was required to reduce inflammatory markers in the tissue after injury in aged mice. However, initiating αIL17 injections at the time of injury stunted infiltration of CD45^+^ immune cells including IL4^+^ cells that are critical for tissue repair, suggesting the importance of acute IL17 during wound healing to attract the immune effectors and initiate tissue regeneration (**Fig. S23b**). We then tested initiation of αIL17 injections one week after injury to allow immune infiltration before immunotherapy treatment (**Fig. 6a**). Using aged Il4^tm1Lky^ mice (4Get), which have a fluorescent reporter for IL4 expression, we found that both αIL17A and αIL17F treatment one week after injury significantly increased the number of IL4^+^ eosinophils and CD4 T cells 3 weeks after injury.

**Figure 6.**
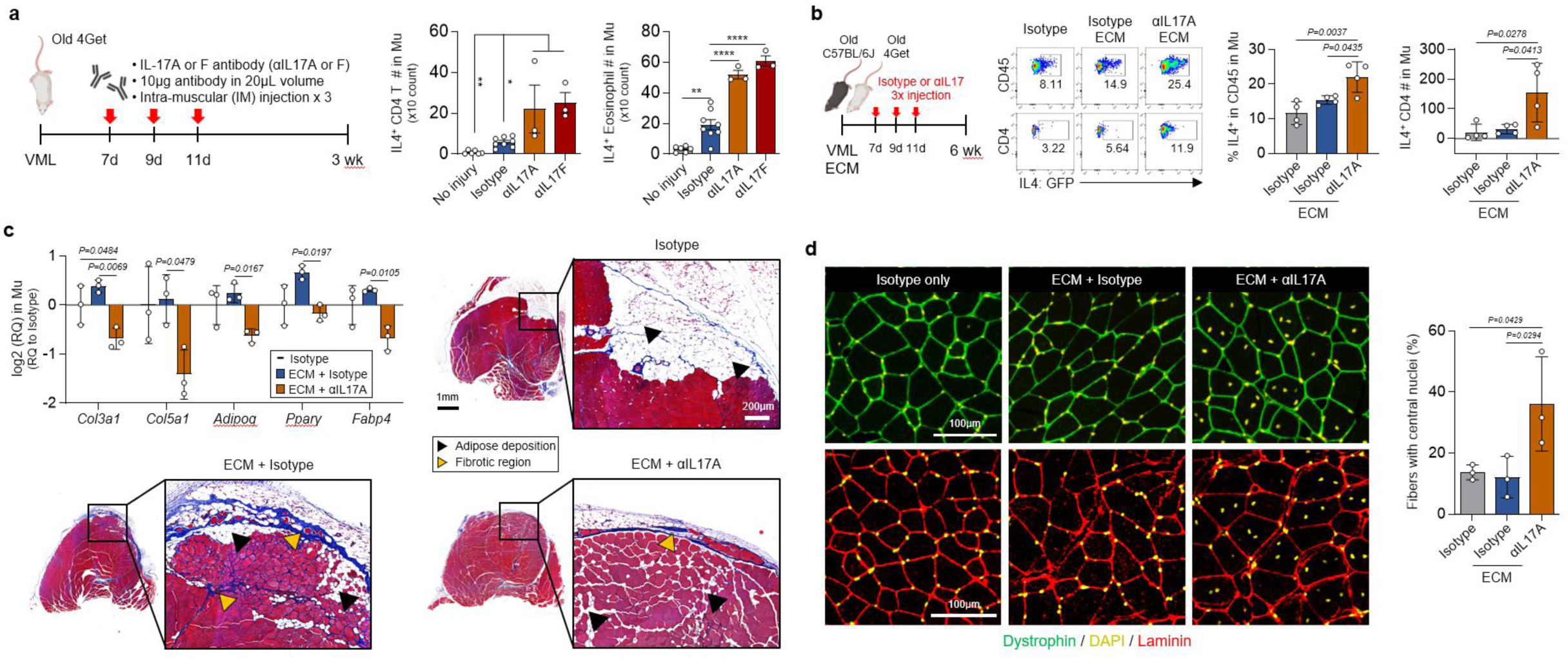
Local IL17 suppression rejuvenates the type 2 immune response to injury and ECM to restore tissue repair and reduce fibrosis in old animals. **a**, Schematic illustration of experimental design (left) and quantification of flow cytometry data for IL4^+^ CD4 T cells and eosinophils in muscle 3 weeks after injury (right). **b**, Experimental schematics (left), and representative images flow cytometry showing IL4^+^ CD45 or CD4 T cells (middle) and quantification of IL4^+^ cell populations in muscle 6 weeks after injury and ECM treatment (right). **c**, Quantification of genes associated with fibrosis or adipogenesis in muscle (top left), and transverse section of the quadricep muscle 6 weeks after injury stained with Masson’s Trichrome. **d**, Immunofluorescence images of the quadricep muscle 6 weeks after injury stained with dystrophin (top) or laminin (bottom). Quantification of muscle fibers with central nuclei are shown (right). Nuclei were stained with DAPI (represented in yellow). One-way ANOVA with Tukey’s multiple comparisons test (a-d). \**p*<0.05, \*\**p*<0.01, and \*\*\*\**p*<0.0001. For all bar graphs, data are mean ± s.e.m (**a**) or s.d. (**b**-**d**).

### Combination therapy of pro-regenerative ECM and αIL17 rejuvenates muscle repair in aged mice

Since αIL17 treatment alone restored IL4 expression after injury in aged animals, we tested a combination therapy approach with pro-regenerative ECM and different types of αIL17 (**Fig. 6b**). Muscle wounds in aged animals received ECM treatment at the time of VML injury and local injection of αIL17 one week after injury. Six weeks after VML in aged 4Get mice, all combination therapies increased the number of CD45^+^ cells in muscle (**Fig. S24**), however, only the αIL17A treated animals showed significant increases in IL4^+^CD45^+^ immune cell percentage and IL4^+^CD4^+^ T cell numbers compared to the ECM alone group, albeit in small numbers at this later time point.

We then explored the therapeutic response to ECM-αIL17 combination in the muscle wound and tissue repair. Six weeks after injury, muscle tissue from aged C57BL/6J mice treated with ECM-αIL17A expressed significantly lower levels of fibrosis- or adipose-associated genes that CoGAPS analysis identified as increasing in aged animals (**Fig. 6c, top left panel**). Specifically, collagen type III a1 (*Col3a1*) and collagen type V a1 (*Col5a1*) decreased significantly only with ECM-αIL17A combination treatment. Additionally, adipose-associated genes, such as *Pparγ, Fabp4*, and *Adipoq* all dramatically decreased with αIL17A treatment. Other fibrosis-related genes, such as *Fap* and *Pdgfa*^57^, decreased with αIL17A or αIL17A + αIL17F (**Fig. S25**). Expression of the inflammatory gene *Mmp13* also significantly decreased with any combination of αIL17 treatments. Differences in gene expression profile between the αIL17A and αIL17F-treated groups suggest that there is a distinction between IL17A and IL17F signaling pathways.

Histological evaluation of the muscle defect in aged animals treated with a combination therapy further supported the gene expression results with increased repair and reduced fibrosis depending on the form of IL17 neutralization (**Fig. 6c** and **Fig. S26**). Aged mice treated with ECM alone demonstrated collagen deposition and fibrosis that decreased with αIL17A or αIL17A + αIL17F treatment. Adipose deposition in the repaired tissue also decreased with ECM-αIL17A combination treatment, supporting the gene expression results. Additionally, a combination treatment induced nuclear repositioning from the periphery to the center of muscle cells (**Fig. 6d**), a characteristic of repairing muscle tissue that did not occur in animals treated with injury or ECM alone. Nuclear positioning is critical in muscle fiber function as the regenerating myofibers after muscle damage can be characterized by centrally localized nuclei^58,59^. Quantification of muscle cells with centrally-located nuclei demonstrated that ECM-αIL17A combination treatment significantly increased the percentage of regenerating muscle fibers compared to the injury or ECM alone groups (**Fig. 6d, right panel**). Altogether, we demonstrate that aging significantly alters the immune and stromal response to muscle injury and therapeutic biomaterials in the local tissue, regional lymph nodes and systemically in blood. These altered responses, characterized by increased IL17, reduced IL4 and excess fibrosis and adipogenesis, can be mitigated with a combination therapy of a pro-regenerative ECM material and an IL17 antibody to promote the repair capacity in aging tissue.

## Discussion

In this work, we uncovered age-related changes associated with type 3 (IL17) immunity that are present in secondary lymphoid organs, that is further exacerbated after muscle injury and treatment with a regenerative ECM biomaterial. Repaired tissue in the aged animals is characterized by excessive fibrosis and adipose tissue with treatment. Single cell analysis revealed excessive collagen activity and abnormalities in antigen presentation with myeloid cells in the aged animals. Cell communication highlighted diminished immune-stromal cell interactions with aging, particularly in the fibroblasts, vascular-related clusters (endo/peri), and T cells, which had an altered secretome. Signaling among the cells also emphasized a unique communication between the senescent fibroblast and T cells, which correlated with activation of Th17-associated TFs in T cells of old animals. Combination therapy of the ECM scaffold with an IL17 neutralizing antibody in the aged animals restored, in part, the pro-regenerative immune response and tissue repair while reducing fibrosis and excess adipose.

Tissue injury mobilizes the immune system and uncovers new age-associated dysfunctions that may not be otherwise apparent. Aging is associated with numerous chronic diseases and increased incidence of cancer^60^. Healthy aging though, even without overt disease, results in longer recovery times from tissue injury. Changes in cellular composition with aging may be in part responsible for reduced healing capacity including decreased endogenous stem cell numbers and activity, in addition to reduced fibroblast heterogeneity^8,9^. However, the pivotal role of the immune system in the response to tissue injury and directing tissue repair is critical to consider as there are many age-related changes in the immune system. Even the epigenetic changes that have been implicated in age-associated repair dysfunction^61^ may extend to the aging immune response to tissue damage as we observed a different secretome of aged Th17-skewed cells cultured in similar conditions to young T cells that is likely due to epigenetic changes. As regenerative medicine strategies are moving to target the immune system, understanding these age-associated immune changes will be critical to develop regenerative immunotherapies that are relevant to the older patient populations that are more likely to suffer from delayed or inadequate tissue repair. Finally, as biological age does not always correlate with chronological age, relevant diagnostics and personalized therapeutic approaches may be needed.

While multiple regenerative medicine therapies are available, we chose ECM biomaterials to evaluate in an aging environment because of their clinical use^16^. Few regenerative medicine therapies have reached standard clinical practice and now even the mechanism of long-studied stem cell therapies is being reconsidered, as non-viable cells appear to promote similar therapeutic responses as viable cells^62^. ECM biomaterials derived from allograft and porcine sources are approved for wound healing and reconstructive surgery applications, orthopedic, and ophthalmologic indications^16,63,64^. ECM materials contain a complex mixture of proteins, proteoglycans, and even matrix-bound vesicles that likely all contribute to damage signals and others as yet determined factors that mobilize multiple immune and stromal cell types to promote tissue repair. It is likely that other regenerative medicine approaches, cellular, growth factors or small molecules, will exhibit similar reduced efficacy with aging that we found using ECM.

Aged animals exhibited a baseline inflammatory state with more CD8^+^ T cells and Th17 cells, the latter being most predominant in the lymph nodes. In the muscle tissue, however, IL17 expression was only observed after injury and ECM treatment, which induced the most significant increases in IL17. While *Il17a* and *f* were observed in the lymph node, only *Il17f* gene expression was found in the muscle tissue. Injury in the older animals uncovered many age-related signatures associated with IL17 and its signaling, and this was further exacerbated with ECM implantation. The cytokine IL17 is a component of the host defense against extracellular pathogens^36,65^, but is also associated with fibrosis and fibrotic disease^55,56^, suggesting a common mechanism of “walling off” uncontrolled pathogens and maintaining barrier surfaces and microbiome balance. While IL17 is important for the recruitment of effector immune cells for wound repair and host defense, its chronic state with aging can further induce carcinogenesis, fibrosis, and inappropriate immune responses. Age-associated commensal dysbiosis may contribute to the excess IL17 in addition to senescence-induced immunomodulation that promotes IL17^66^. As mice are reared in a controlled lab environment, the increased aged-associated IL17 related to gut dysbiosis may be even greater and more variable in people that have more diverse environment exposure, diet, and etc.

Intercellular communication analysis by Domino uncovered active immune-stromal module interactions in young animals that were impaired and limited in an aging environment. Aging notably changed the signaling from the fibroblasts, especially the senescent fibroblasts, and demonstrated a unique communication with the T cells, which further provided a biological mechanistic of type 3 adaptive immune skewing in old animals. Young mice demonstrated active immune-stromal communication associated with vascular development (signaling with Endo/Peri cluster), which is well recognized for its roles in tissue repair. Interestingly, the aged Th17-skewed cells secreted more VEGF *in vitro* compared to the young T cells, suggesting a possible epigenetic memory associated with vascular insufficiency. In addition to VEGF, the aged Th17 cells also secreted increased levels of LIF, a component of the Stat3 and Wnt signaling pathway involved in vascular development. Vascular insufficiency and impaired VEGF signaling is a hallmark of aging, particularly in the microvasculature which is a necessary component of tissue repair regeneration^67^. We also previously showed that inflammatory (Th17)-induced senescent cells produce a SASP that upregulated Wnt signaling genes, such as Wnt5b and Wnt11, further implicating Wnt signaling in age-related repair dysfunction.

In summary, the immune system represents an exciting new therapeutic target for regenerative medicine. However, the complexity of the immune system in people and variability related to intrinsic genetic, sex differences, exposure history and environmental factors that only increases with age must be considered in therapeutic design. Combination therapies, a standard approach in cancer treatment, should be extended to regenerative medicine where complex interactions between the immune system, stem cells, and the vascular system contribute to repair outcomes.

## Acknowledgements

This work was funded in part by the National Institutes of Health (NIH) Pioneer Award DP1AR076959 (J.H.E.), Bloomberg∼Kimmel Institute, Morton Goldberg Professorship (J.H.E.), the Department of Defense, R01EB028796 (J.H.E.), R01AG063801(J.H.E.). A.R. and D.R.M. were funded through NSF GRFP DGE-1746891. We thank F. Housseau for providing the IL17A-GFP mice. Schematic illustrations created with BioRender.com.

## Author contributions

J.H., and J.H.E. conceptualized the study, drafted, and edited the figures and manuscript. J.H., J.H.E., and D.M.P. contributed to experimental design and interpretation of the results. J.H. performed animal surgery model and analyzed experimental results (flow cytometry, qRT-PCR, proteome profiler, histology, and analysis). C.C. wrote software and assisted in single cell RNA sequencing analysis and computational methodology. K.K. assisted with computational methodology. A.R., J.C.M., and D.R.M. assisted with animal breeding and spectral flow cytometry analysis. H.H.N. contributed to conducting histological evaluation. A.N.P, B.Y., N.R., E.G.G., and E.M.M. assisted with *in vitro* T cell culture or subsequent genomic profile assessment. E.J.F. advised in the bioinformatics analysis. F.H., S.G., and A.J.T. assisted with generalized gene expression interpretation and flow cytometry results.

