## Supplementary information for "Age-associated Senescent - T Cell Signaling Promotes Type 3 Immunity that Inhibits Regenerative Response"

**Materials and Methods**

Surgical procedures and implantation

All animal procedures were approved by Johns Hopkins University Institutional Animal Care and Use Committee protocol. Mice aged 6 week (young) or 72 week (aged) were obtained from the Jackson Laboratory (C57BL/6J: stock #00064). 4Get mice (stock #004190) were obtained from the Jackson Laboratory and bred in-house. IL17A-GFP mice (courtesy of F. Housseau, Johns Hopkins, MD) were bred in-house. The bilateral muscle defects in quadricep were created as previously described[1]. The defects were either filled with 0.05 cc of 200mg/ml biomaterial scaffold. Decellularized porcine extracellular matrix (ECM) was used as a biological scaffold in 0.05 ml at a concentration of 200 mg/ml in phosphate-buffered saline (PBS). Control surgeries were treated with 0.05 ml of PBS. All materials were sterilized with UV before use. Immediately after surgery, mice were given subcutaneous injection of carprofen (Rimadyl, Zoetis) at 5 mg/kg for pain relief. For analysis, mice were euthanized at 1, 3, or 6 weeks after surgery, and various tissues (blood, inguinal lymph node, or muscle) were extracted. All animal procedures in this study were conducted in accordance with an approved Johns Hopkins University IACUC protocol.

Tissue ECM Preparation

Porcine-derived tissues (Wagner Meats, Mt. Airy, MD) were processed following a protocol previously described[1]. Tissues were formulated into a paste with particle sizes no larger than 5 $mm^2$ and rinsed thoroughly with distilled water. Tissues were then incubated in 3 % peracetic acid (Sigma) on a shaker at $37^oC$ for 4 hours. pH was adjusted to 7 with running distilled water and PBS rinsing, and tested after solution was freshly changed. Samples were then transferred to a 1 % Triton-X100 (Sigma) + 2 mM sodium EDTA (Sigma) solution on a stir plate at 400 rpm, room temperature for 3 days. Tissues were then rinsed thoroughly with distilled water and incubated in 600 U/ml DNase I (Roche Diagnostics) for 24 hours. Tissues were rinsed with distilled water, frozen at $-80^oC$ and lyophilized for at least 3 days. Finally, dry sample was turned into a particulate form using a SPEX SamplePrep Freezer/Mill (SPEX CertiPrep). ECM powder was stored in $-20^oC$ until use, and UV sterilized immediately before use.

Non-surgical animal experiments

For experiments comparing base line immunological difference between young and aged, no surgery was given to young or aged animals, and blood or inguinal lymph node was analyzed using flow cytometry, qRT-PCR, proteome profiler and histological evaluation. Cytokine expression in blood was analyzed using proteome profiler cytokine array (R&D systems) according to the manufacturer's directions.

qRT-PCR

For total mRNA expression in muscle and inguinal lymph node, lysis was conducted on whole tissues using TRIzol at 1 or 6 week after surgery. RNA purification was performed using RNeasy Plus Mini kit (Qiagen). PCR was all performed using TaqMan Gene Expression Master Mix (Applied Biosystems) according to the manufacturer's directions. Briefly, 2 ug of mRNA was synthesized into complementary DNA (cDNA) using Superscript IV VILO Master Mix (Thermo Fisher Scientific), and was used at 100ng/well in a total volume of 20μl of PCR. All qRT-PCRs were performed on the StepOnePlus Real-Time PCR System (Thermo Fisher Scientific). Rer1, OAZ1, and Hprt were used as the reference gene and experimental groups were normalized to either no surgery or saline-treated controls. Low-expressing mRNA transcripts were pre-amplified using the TaqMan Pre-Amp System (Thermo Fisher Scientific) following manufacturer's recommendations with 10 cycles of amplification.

Flow Cytometry

Whole muscle or inguinal lymph node was harvested either without injury, 1, 3, or 6 weeks after surgery. Muscle tissues were obtained by cutting the quadriceps from the hip to the knee, finely diced and digested for 45 min at 37$^{o}$C with 1.67 Wünsch U/ml Liberase TL (Roche Diagnostics) and DNaseI (0.2 mg/ml; RocheDiagnostics) in RPMI 1640 medium (Gibco). The digested tissues were ground through 70 µm cell strainers (Thermo Fisher Scientific) and washed multiple times with PBS. For intracellular staining, cells were stimulated for 4 hrs with Cell Stimulation Cocktail plus protein transport inhibitors (eBioscience) diluted in RPMI 1640 medium supplemented with 10 % fetal bovine serum (FBS). Cells were then washed and surface-stained, followed by fixation/permeabilization (Cytofix-Cytoperm, BD) and intracellular markers. Flow cytometry was performed using Attune NxT Flow Cytometer (Thermo Fisher Scientific) or Cytek Aurora (Cytek). Cells were stained with the antibody panels listed in Supplementary Table S1.

Splenocyte isolation and Th17 differentiation *in vitro*

Spleens from young or aged mice were ground through 70 µm cell strainers (Thermo Fisher Scientific) and washed multiple times with PBS. Cells were then incubated with ACK lysing buffer (Thermo Fisher Scientific) in dark for 10 minutes for red blood cell lysis, followed by multiple PBS wash. Naïve CD4$^{+}$ T cells were then isolated using Naïve CD4$^{+}$ T Cell Isolation Kit (Miltenyi Biotec). The cells were then differentiated using CellXVivo mouse Th17 differentiation kit (R&D systems) for 5 days. For flow cytometry analysis, the cells were collected and stained for flow cytometry. For proteome analysis, the media was changed to T cell culture media (RPMI 1640 with 10% FBS, 1% Penicillin-Streptomycin, 1mM Sodium Pyruvate, 10mM HEPES, and 50nM 2-Mercaptoethanol) on day 5, then cultured for additional 2 days. Supernatant was analyzed using proteome profiler cytokine array (R&D systems) according to the manufacturer's directions.

Coculture of splenocytes with senescent fibroblast

Spleens from young or aged mice were ground through 70 µm cell strainers (Thermo Fisher Scientific) and washed multiple times with PBS. Cells were then incubated with ACK lysing buffer (Thermo Fisher Scientific) in dark for 10 minutes for red blood cell lysis, followed by multiple PBS wash. CD4$^{+}$ T cells were then isolated using CD4$^{+}$ T Cell Isolation Kit (Miltenyi Biotec). Murine dermal fibroblasts were isolated from 5 week old female mice. Mice were euthanized, shaved, and a 1x2cm section of dermal skin was removed and cut into small pieces before digestion. The dermal sections were digested for 1 hour at 37C in 0.5 mg/ml Liberase TM (Roche Diagnostics) in serum free RPMI 1640 shaking at ~150rpm. The digestion was neutralized with complete RPMI 1640 supplemented with 10% fetal bovine serum, 1% penicillin/streptomycin, 1x sodium pyruvate, and the digested skin was pelleted *via* centrifugation. The digested skin was resuspended in complete RPMI 1640, divided into 3 T-175 flasks, and incubated at 37C, 5% CO2 undisturbed for 4 days. On day 4, the cells were washed with PBS and the culture media was changed to complete MEM supplemented with 10% fetal bovine serum, 1% penicillin/streptomycin to promote only fibroblast survival. Once 80% confluent, the fibroblasts were placed in a minimum amount of media and an Xstrahl CIXD X-ray irradiator was used to deliver a dose of 10 Gy to the cells. The irradiated cells were washed with PBS and fresh complete MEM was added. The irradiated cells were incubated at 37C 5%CO2 for 10 days to allow for the senescent phenotype to develop ubiquitously. Control quiescent fibroblasts were obtained by culturing fibroblasts in low-serum MEM supplemented with 0.5% fetal bovine serum, 1% penicillin/streptomycin. Fibroblasts and $CD4^+$ T cells were cocultured using Transwell (Corning®, 100,000 quiescent or senescent fibroblasts in the insert, 500,000 T cells in the well) for 5 days.

IL17 neutralization treatment

Mice received 3 injections 20 µl intra-muscular injections of isotype control (rat IgG2a, R&D systems), anti-IL17a (100 µg/ml, R&D systems), anti-IL17f (100 µg/ml, R&D systems), or anti-IL17a and anti-IL17f combined, every other day. All mice received treatments either at the day of surgery (for dosing experiment) or at 1 week after surgery (all other experiments), and were harvested at 3 or 6 weeks after surgery.

NanoString gene expression analysis

Inguinal lymph nodes from no treat mice were used to isolate mRNA for NanoString analysis. Gene expression was evaluated using the NanoString AutoImmune Profiling Panel (NanoString Technologies, Inc.). 100 ng of RNA was added to a probe-set mixture, and hybridized for 20 hours at 65°C. All samples were processed using a NanoString Prep Station under high sensitivity mode, and mRNA target transcripts were counted using the nCounter digital analyzer system (NanoString Technologies, Inc.). Data was analyzed using nSolver software.

Histopathology

Tissues were harvested 1 or 6 weeks after surgery and fixed in 10 % neutral buffered formalin for 48 hours. Tissues then underwent stepwise dehydration in EtOH, followed by xylenes, and embedded in paraffin. Tissue samples were sectioned as 6 µm slices, then stained for histopathological examination using Masson's Trichrome, hematoxylin and eosin, or immunofluorescence. Dystrophin and Laminin were stained using tyramide signal amplification method with Opal-570 (PerkinElmer, catalog no. FP1488001KT). Briefly, after blocking with bovine serum albumin for 1 hours, the primary antibody was incubated at room temperature for 30 min, followed by 10 min of incubation with horseradish peroxidase (HRP) polymer-conjugated secondary antibody, and 10 min of Opal. Slides were then counterstained with 4′,6-diamidino-2-phenylindole (DAPI) for 5 min before being mounted using DAKO mounting medium (Agilent, catalog no. S302380-2). Imaging of the histological samples was performed on a Zeiss Axio Imager A2 and Zeiss AxioVision software version 4.2. Immunofluorescent images were analyzed using ImageJ software.

Collection of single cell data sets

Drop-seq, a single cell microfluidics encapsulation technique, was used to prepare libraries for $CD45^+$ enriched cell populations isolated from mouse quadriceps 1 week after the treatments. For the CD45+ enriched populations, dead cells were removed using the Miltenyi Biotec Dead Cell Removal Kit followed by Miltenyi Biotec CD45 MicroBeads to separate $CD45^+$ and $CD45^-$ cells. After separation, an equal amount of $CD45^+$ and $CD45^-$ cells were pooled directly prior to input to Drop-seq. Drop-seq was run following the McCarroll Lab's December 2015 iteration of their published protocol available from their website (http://mccarrolllab.org/dropseq/).

Data preprocessing and batch effect correction

Seurat was used for most processing steps where other software is not specified[2]. All cell counts were pruned of cells with UMI counts below 200, cells with more than 10% mitochondrial genes, and genes expressed in fewer than 0.1% of cells. We then normalized and scaled the data with regression on UMI count, G2M score, S score and percent mitochondrial genes and integrated the data with Seurat. We then calculated principle components using the top 2000 most variable genes. UMAP and shared nearest neighbor graph construction with subsequent Louvain clustering was then run on principle components.

Cluster composition by condition

To assess cluster contribution, clusters from $CD45^+$ and $CD45^-$ cells were normalized separately to avoid slight differences in percent of $CD45^+$ cells from enrichment by sample skewing normalization. For each sample, total number of cells by cluster were calculated and then normalized to the total of $CD45^+$ or $CD45^-$ cells in the dataset for the sample. The proportions of each sample were then averaged by condition to determine a condition-level average.

Phenotypic assignment of clusters

Seurat's CellCycleScoring function was used to score cells based on expression of a subset of genes previously identified as associated with the G2M or S phase[3]. Differential expression testing for clusters was run using Mann-Whitney U tests. Each cluster was compared against all other clusters. The resulting gene expression profiles were examined to determine cluster phenotype. In many cases, unique expression of marker genes was sufficient to determine cluster identity.

Intercellular signaling networks

We used Domino to investigate potential signaling patterns between clusters of cells. Domino predicts activated transcription factors by cell using SCENIC[4] and then constructs a network connecting transcription factors, receptors, and ligands based on similar expression patterns. Default parameters for network construction were used. Networks were calculated for old and young samples individually and then compared to determine signaling components specific to each condition.

CoGAPS analysis

scCoGAPS[5] was used to perform non-negative matrix factorization (NMF) to identify 10 underlying patterns. Prior to NMF, mitochondrial and ribosomal genes were removed. It decomposes the data two matrices containing sets of low dimensional features, one of which called the pattern matrix contains a set of weights for each cell and the other called the amplitude matrix the corresponding weight for which each gene. The gene weights in the columns of the amplitude matrix indicate how much a gene contributes to the expression pattern identified by the corresponding row of the pattern matrix. The cell weights in the rows of the pattern matrix indicate how strongly a cell is enriched for the feature and can be used to identify cells with similar expression patterns to the feature. Finally, other cell labels (condition of origin, cluster label, etc.) can be compared with feature scores to identify how feature expressions change with respect to experimental variables such that even if there are no differences in cell clustering with ECM treatment or age, gene signatures can still vary significantly and provide functional insights. The resulting amplitude and pattern matrices were subsequently used to identify cells enriched by pattern and the genes driving patterns. Feature numbers were selected to maximize distinct expression signatures.

Statistical Analysis

All analyses of qRT-PCR data used Livak method, where ΔΔCt values were calculated and reported as relative quantification values calculated by 2-ΔΔCt. Data are displayed as mean ± s.d. Statistical analysis was performed using a one-way or two-way ANOVA with Tukey's corrections applied using GraphPad Prism v8, with statistical significance designated at $p \leq 0.05$. All groups were compared to each other for multiple comparisons unless otherwise stated.

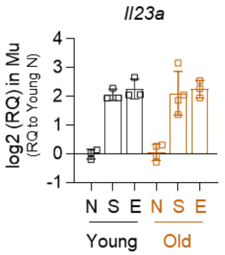


- Young (6 wk), Old (72 wk)
  - ➢ No injury (N)
  - ➢ Injury + Saline (S)
  - ➢ Injury + ECM (E)

| Tukey's multiple comparison | | p |
|---|---|---|
| Young N | Young S | *** |
| | Young E | *** |
| | Old N | NS |
| | Old S | *** |
| | Old E | *** |
| Young S | Young E | NS |
| | Old N | *** |
| | Old S | NS |
| | Old E | NS |
| Young E | Old N | *** |
| | Old S | NS |
| | Old E | NS |
| Old N | Old S | **** |
| | Old E | *** |
| Old S | Old E | NS |

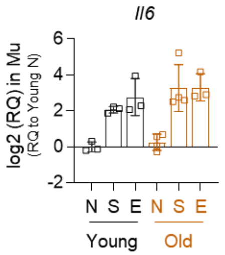


| Tukey's multiple comparison | | p |
|---|---|---|
| Young N | Young S | NS |
| | Young E | ** |
| | Old N | NS |
| | Old S | ** |
| | Old E | ** |
| Young S | Young E | NS |
| | Old N | NS |
| | Old S | NS |
| | Old E | NS |
| Young E | Old N | * |
| | Old S | NS |
| | Old E | NS |
| Old N | Old S | ** |
| | Old E | ** |
| Old S | Old E | NS |

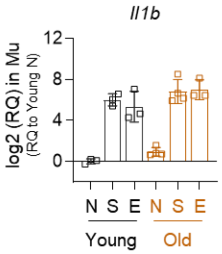


| Tukey's multiple comparison | | p |
|---|---|---|
| Young N | Young S | **** |
| | Young E | **** |
| | Old N | NS |
| | Old S | **** |
| | Old E | **** |
| Young S | Young E | NS |
| | Old N | **** |
| | Old S | NS |
| | Old E | NS |
| Young E | Old N | *** |
| | Old S | NS |
| | Old E | NS |
| Old N | Old S | **** |
| | Old E | **** |
| Old S | Old E | NS |

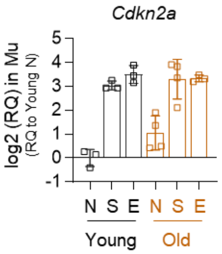


| Tukey's multiple comparison | | p |
|---|---|---|
| Young N | Young S | *** |
| | Young E | **** |
| | Old N | NS |
| | Old S | **** |
| | Old E | **** |
| Young S | Young E | NS |
| | Old N | ** |
| | Old S | NS |
| | Old E | NS |
| Young E | Old N | *** |
| | Old S | NS |
| | Old E | NS |
| Old N | Old S | *** |
| | Old E | ** |
| Old S | Old E | NS |

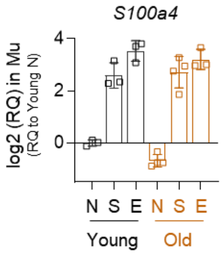


| Tukey's multiple comparison | | p |
|---|---|---|
| Young N | Young S | **** |
| | Young E | **** |
| | Old N | NS |
| | Old S | **** |
| | Old E | **** |
| Young S | Young E | NS |
| | Old N | **** |
| | Old S | NS |
| | Old E | NS |
| Young E | Old N | **** |
| | Old S | NS |
| | Old E | NS |
| Old N | Old S | **** |
| | Old E | **** |
| Old S | Old E | NS |

**Supplementary figure 1**

Quantification of inflammation- or senescence-associated genes in muscle 1 week after injury or treatment. Statistical analysis was performed using a two-way ANOVA with Tukey's multiple comparisons test (n=3-4). For all bar graphs, data are mean ± s.d.

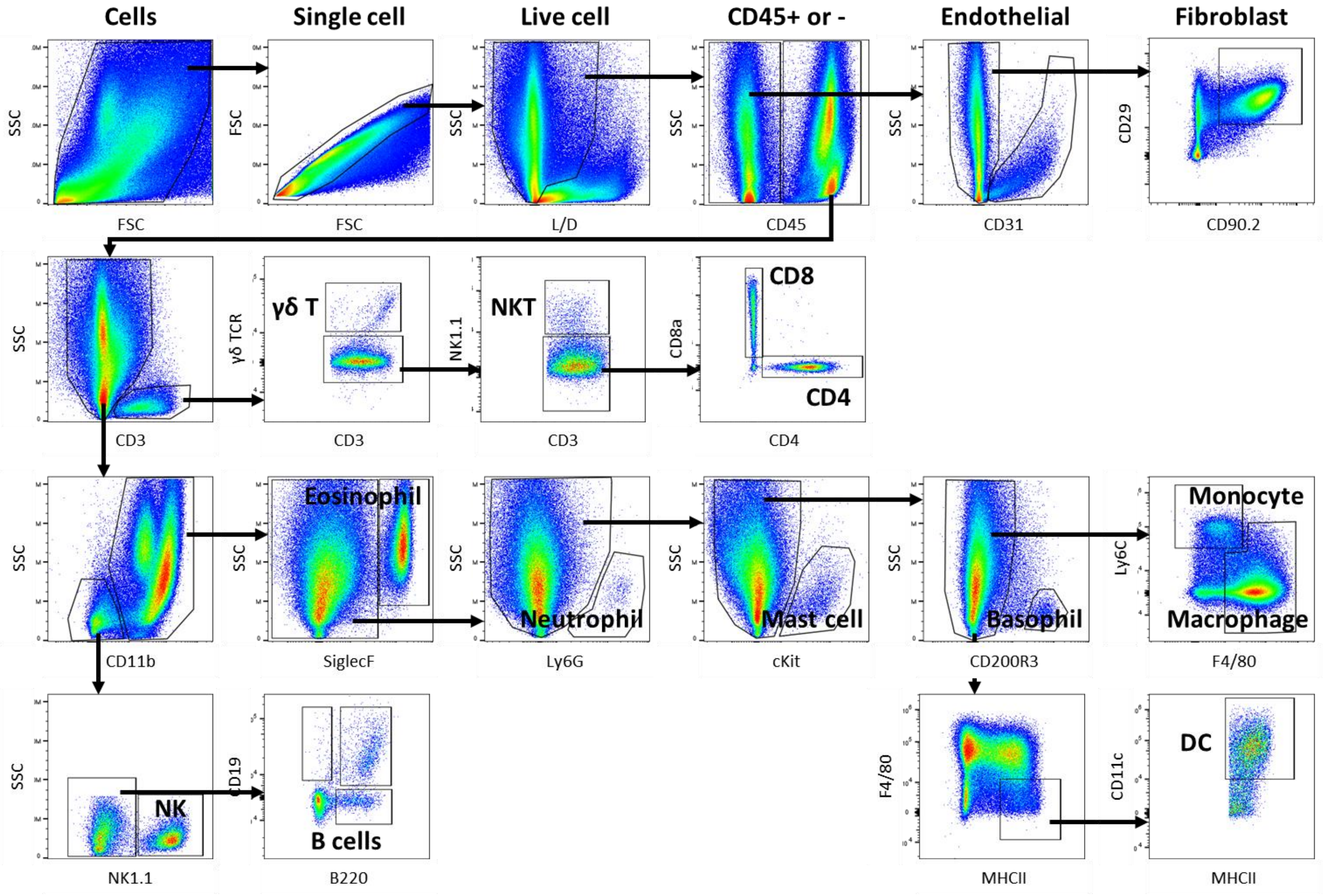


**Supplementary figure 2**

Schematic representation of the gating strategy used to identify indicated cell phenotypes from single cell suspensions from mouse muscle tissue using spectral flow cytometry.

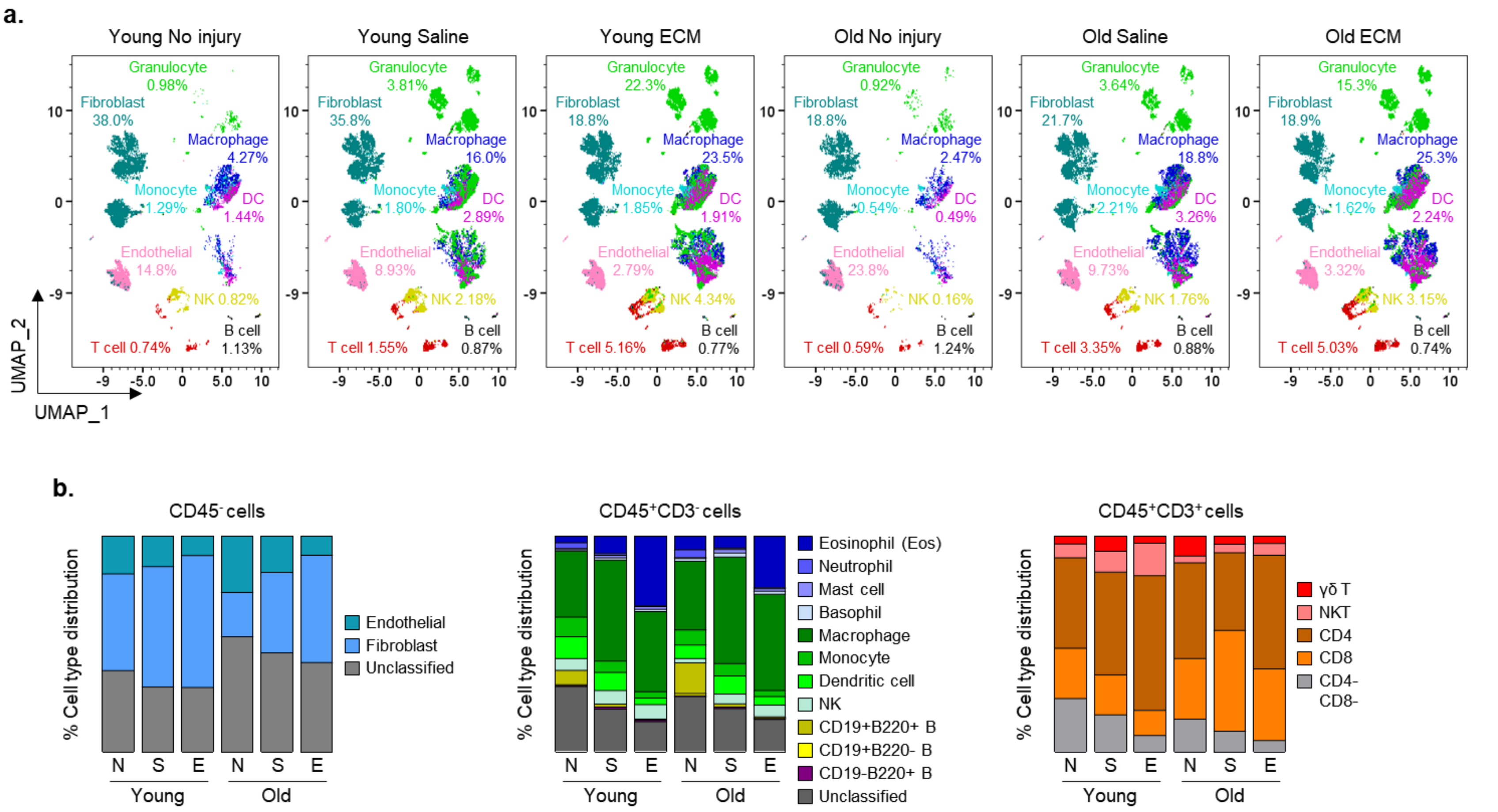


**Supplementary figure 3**

Changes in immune and stromal cell phenotypes 1 week after injury or ECM treatment between young and aged animals. **a**, Overview of UMAP plots from total live cells in muscle showing the population of the indicated cell phenotypes. **b**, Quantification of immune-stromal cells in muscle after treatment as determined by flowcytometry. (**a, b**) Indicated cell population represents average value of n=5 per group.

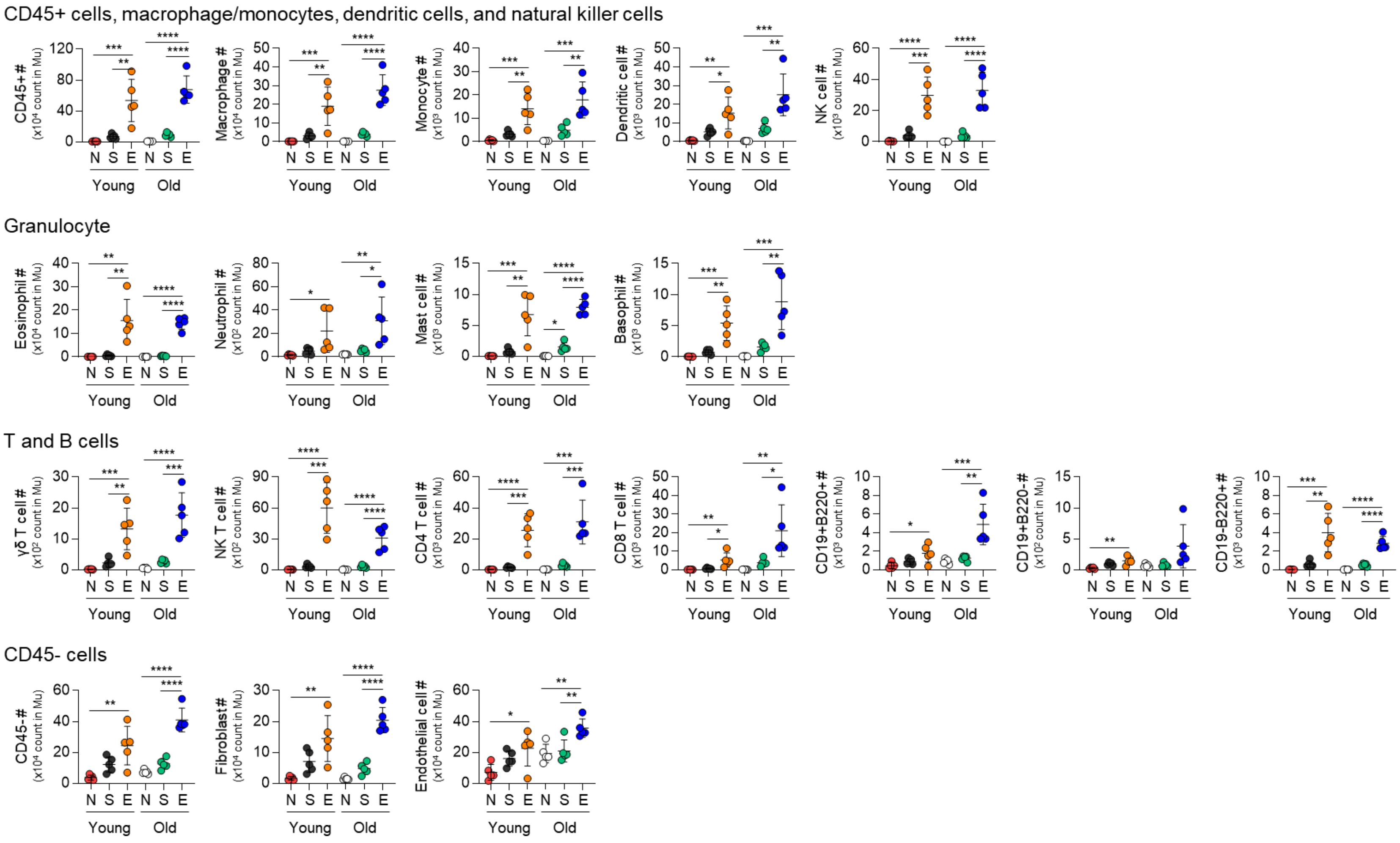


**Supplementary figure 4**

Flow cytometry quantification (cell counts) of the indicated immune and stromal cell phenotypes in muscle from young and aged animals 1 week after injury or treatment. Statistical analysis was performed using a one-way ANOVA with Tukey's multiple comparisons test within the respective age groups (n=5). **p*<0.05, ***p*<0.01, ****p*<0.001, and *****p*<0.0001. For all bar graphs, data are mean ± s.d.

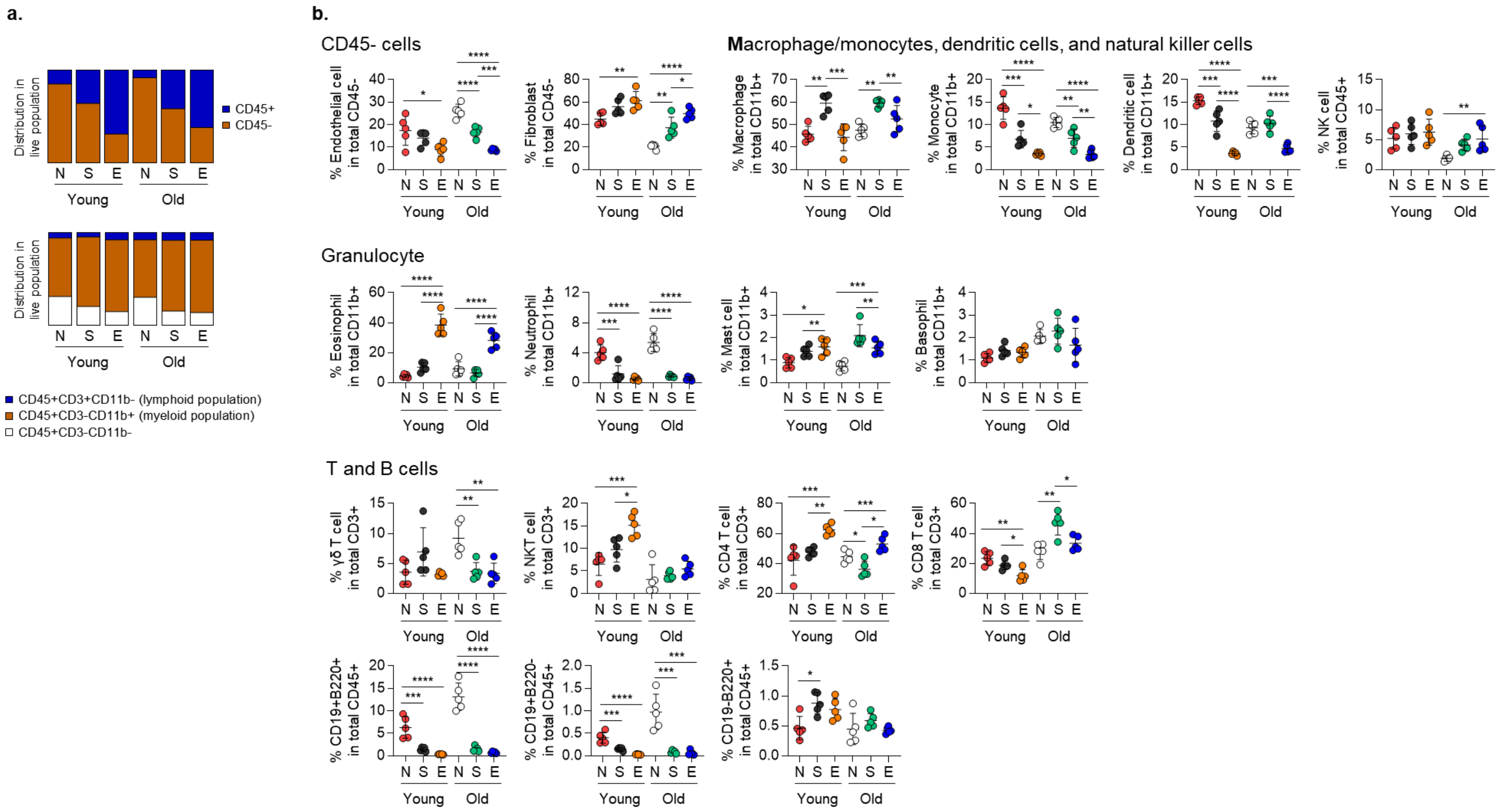


**Supplementary figure 5**

Flow cytometry quantification (cell percentages) of **a,** CD45+/CD45- cells or myeloid/lymphoid cells (indicated cell population represents average value of n=5 per group), and **b,** indicated immune/stromal cell phenotypes in muscle between young and aged animals 1 week after the treatments. Statistical analysis was performed using a one-way ANOVA with Tukey's multiple comparisons test within the respective age groups (n=5). **p*<0.05, ***p*<0.01, ****p*<0.001, and *****p*<0.0001. For all bar graphs, data are mean ± s.d.

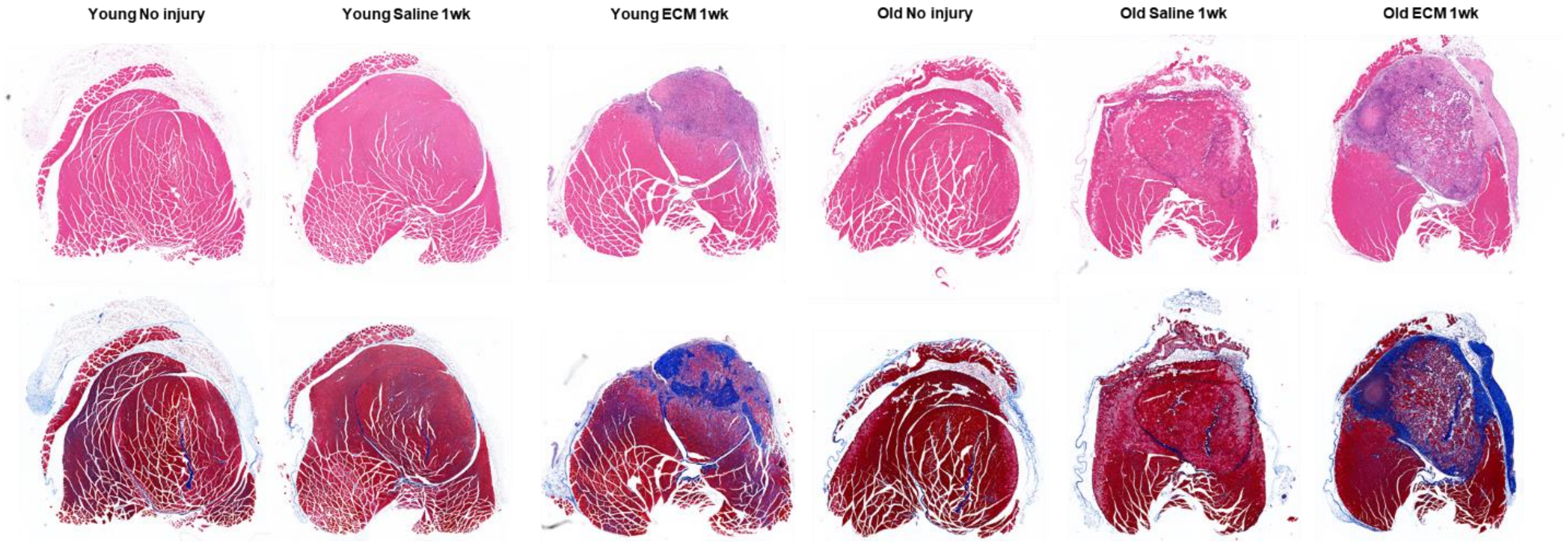


**Supplementary figure 6**

Whole image of transverse section of the quadricep muscle 1 week after injury or ECM treatment stained with H&E (top) or Masson's Trichrome (bottom).

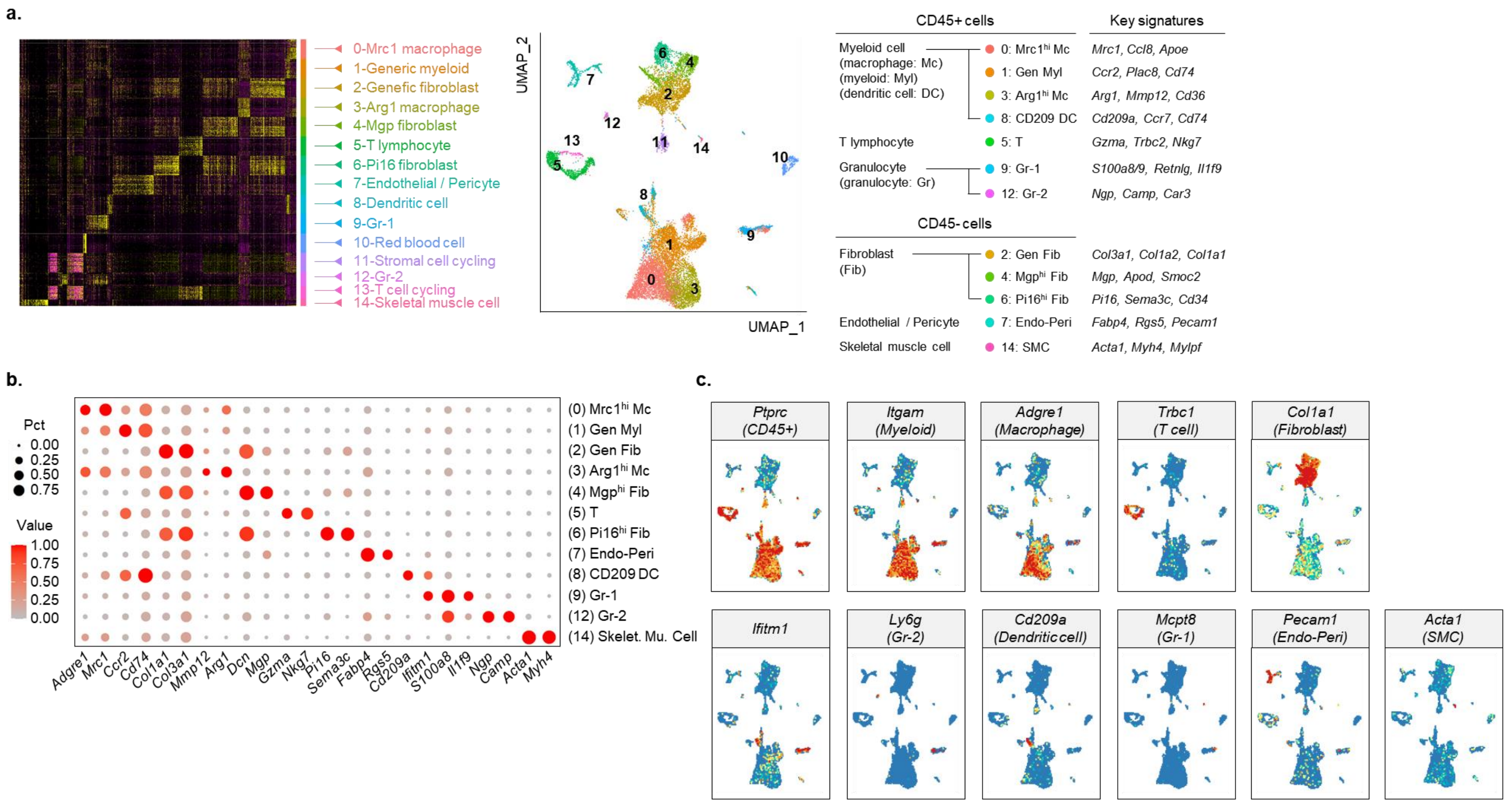


**Supplementary figure 7**

Single cell RNA sequencing-based cell clustering information in young and aged muscle tissue 1 week after injury or ECM treatment. **a**, Heatmap of differentially expressed genes with highest log fold-change from each cluster (left) and UMAP plot with cluster labels and key signature genes (right). **b**, Signature gene markers for single cell clusters. Each dot shows the expression of genes associated with cluster identity. Gene expression after normalization to the maximum averaged expression are shown. **c**, Overview of cell clusters identified in UMAP plots based on the key gene expression.

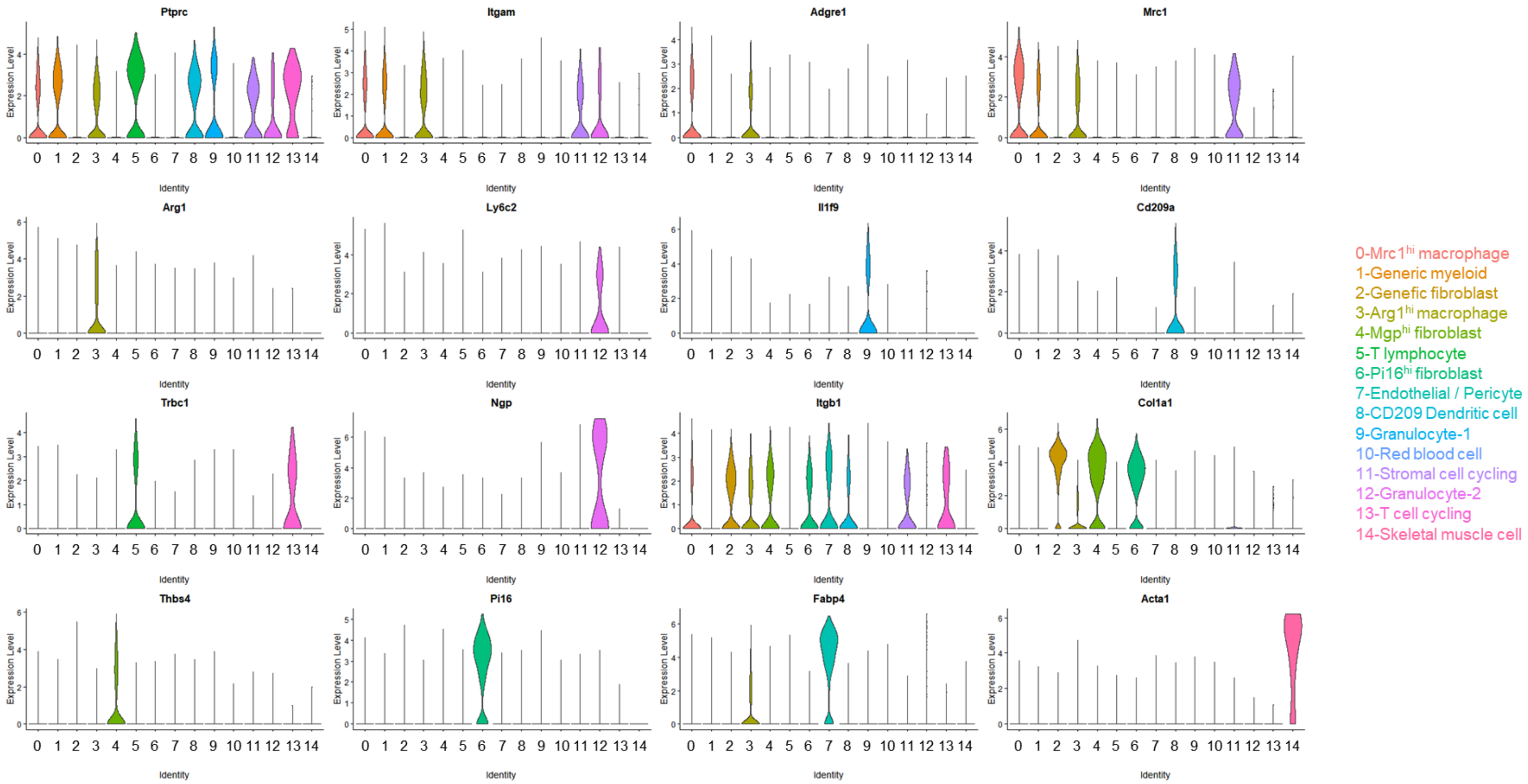


**Supplementary figure 8**

Gene expressions of cluster-defining markers shown in violin plots.

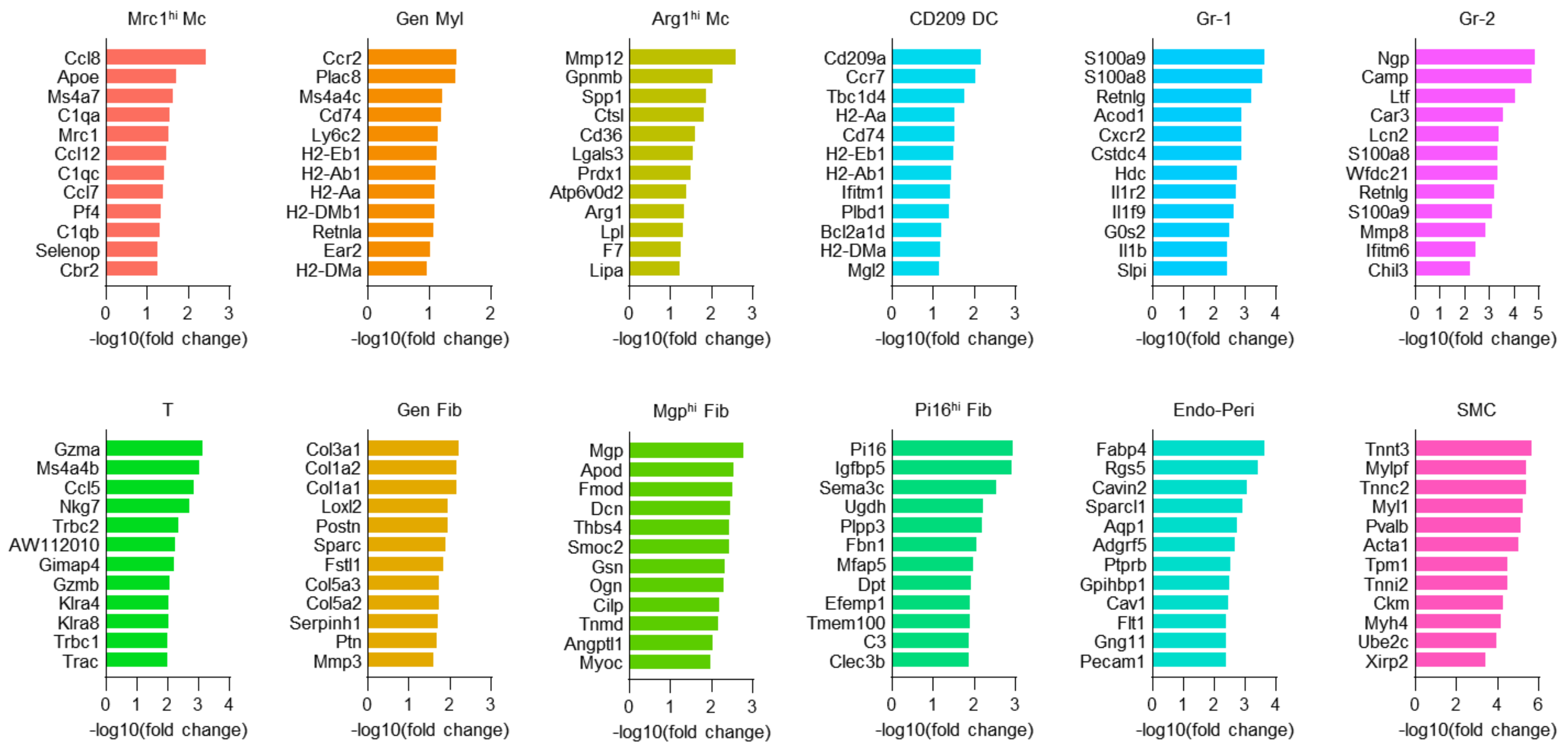


**Supplementary figure 9**

Twelve top differentiated genes of each cluster population. Values represent log10 fold change between each cluster compared to all other clusters.

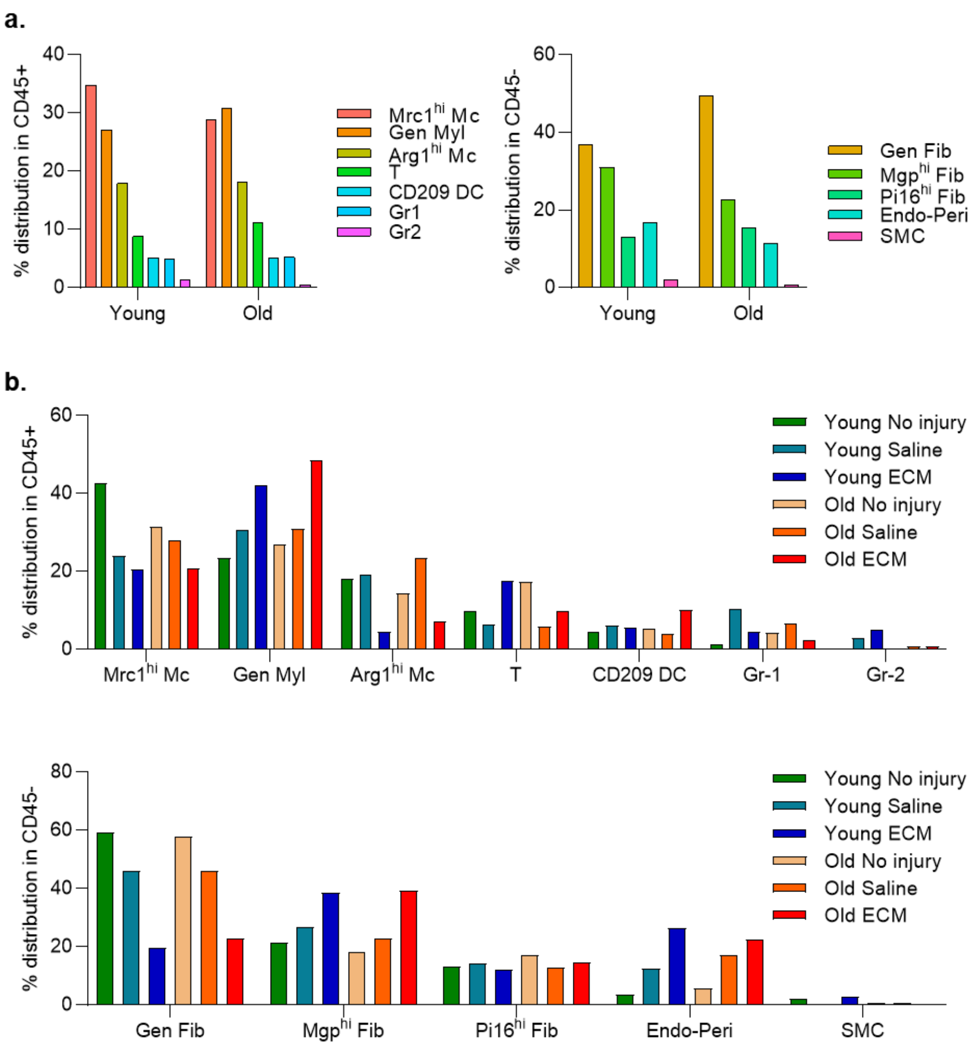


**Supplementary figure 10**

Quantification of each cell cluster between the **a,** age groups or **b,** treatment groups using single cell RNA sequencing dataset**.**

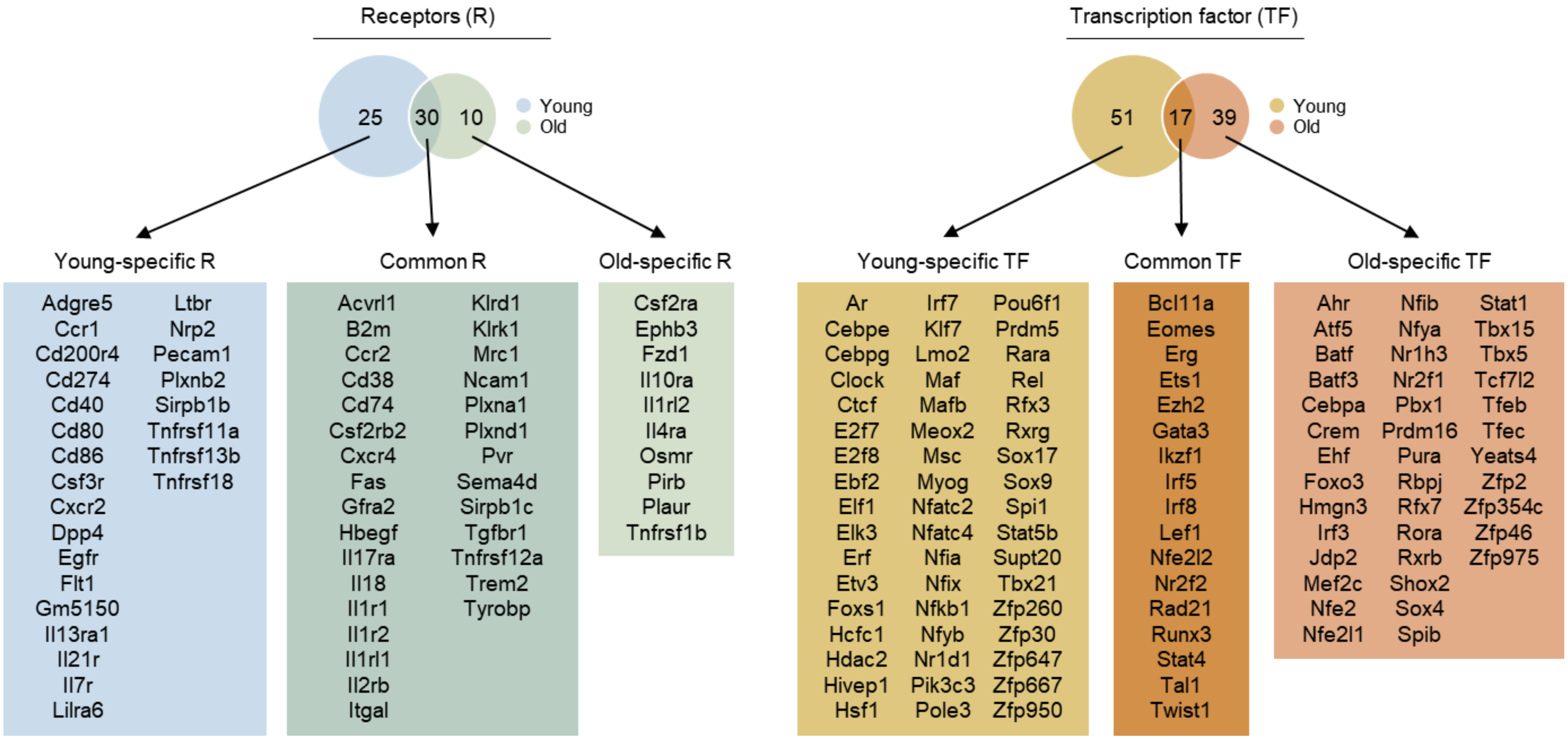


**Supplementary figure 11**

A full list of receptors and transcription factors that are common or specific to young or aged animals using Domino.

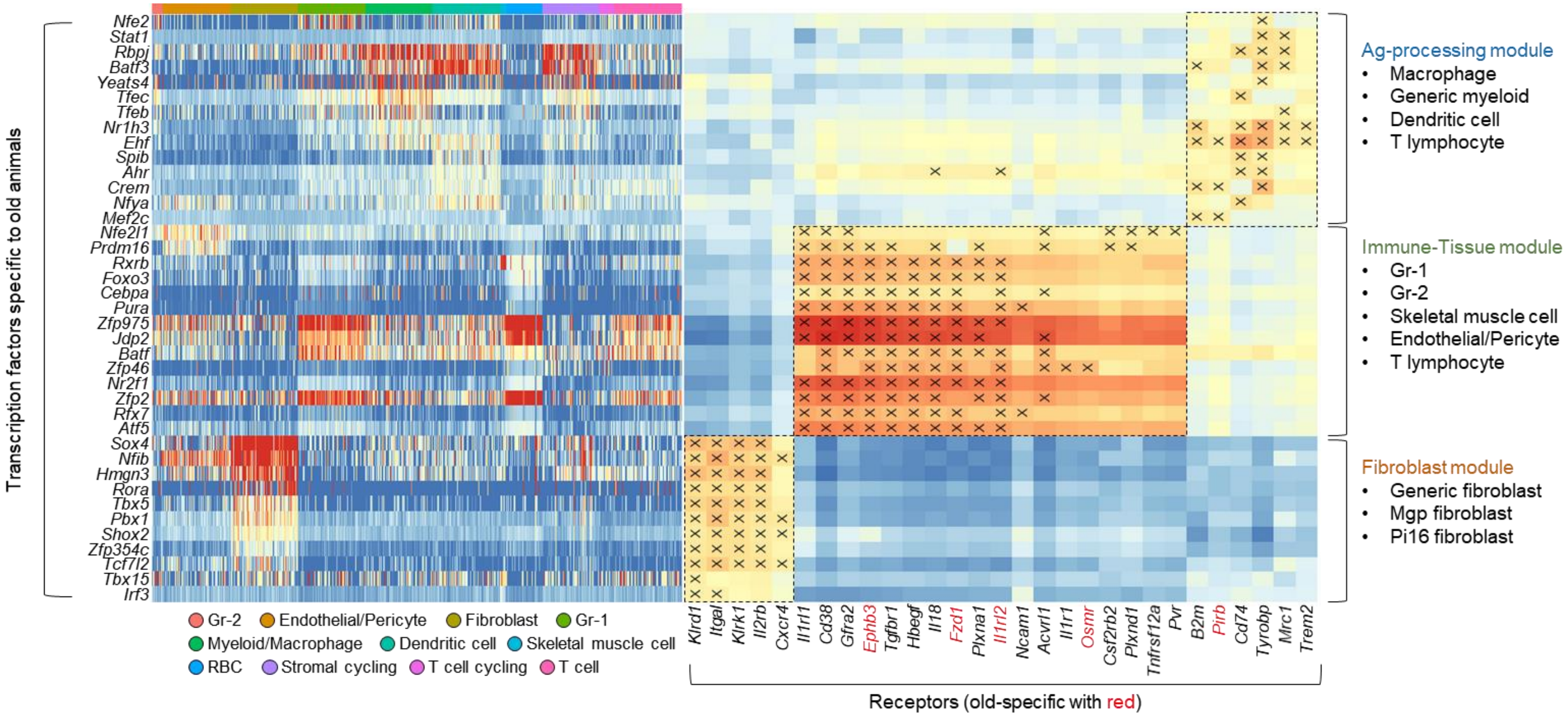


**Supplementary figure 12**

Heatmaps of activation score for aged animal-specific transcription factors and their correlation with receptor expression. Transcription factors specific to aged animals are labelled on the right, and the correlated receptors are labeled on top right.

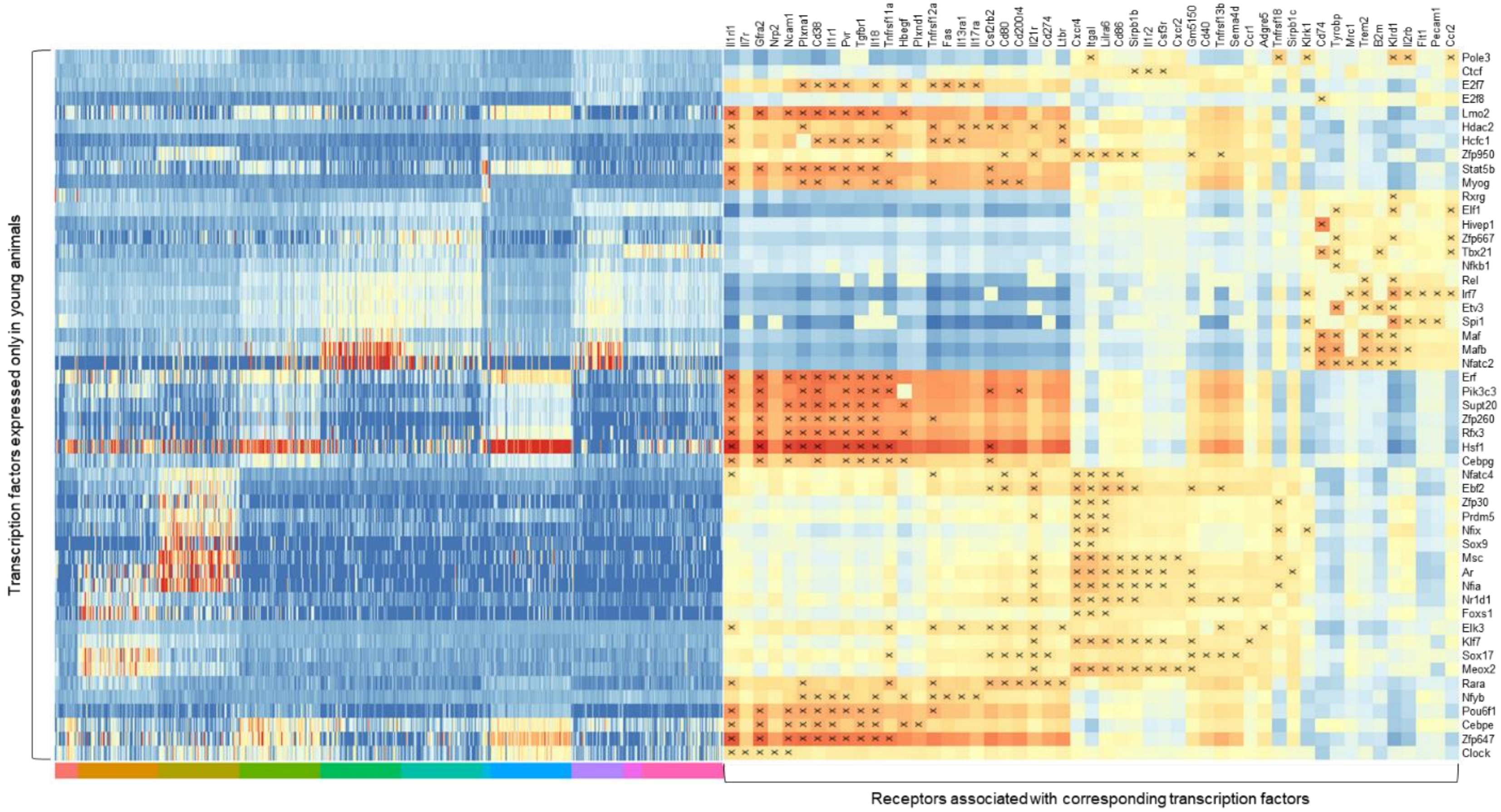


**Supplementary figure 13**

Heatmaps of activation score for young animal-specific transcription factors and their correlation with receptor expression. Transcription factors specific to young animals are labelled on the right, and the correlated receptors are labeled on top right.

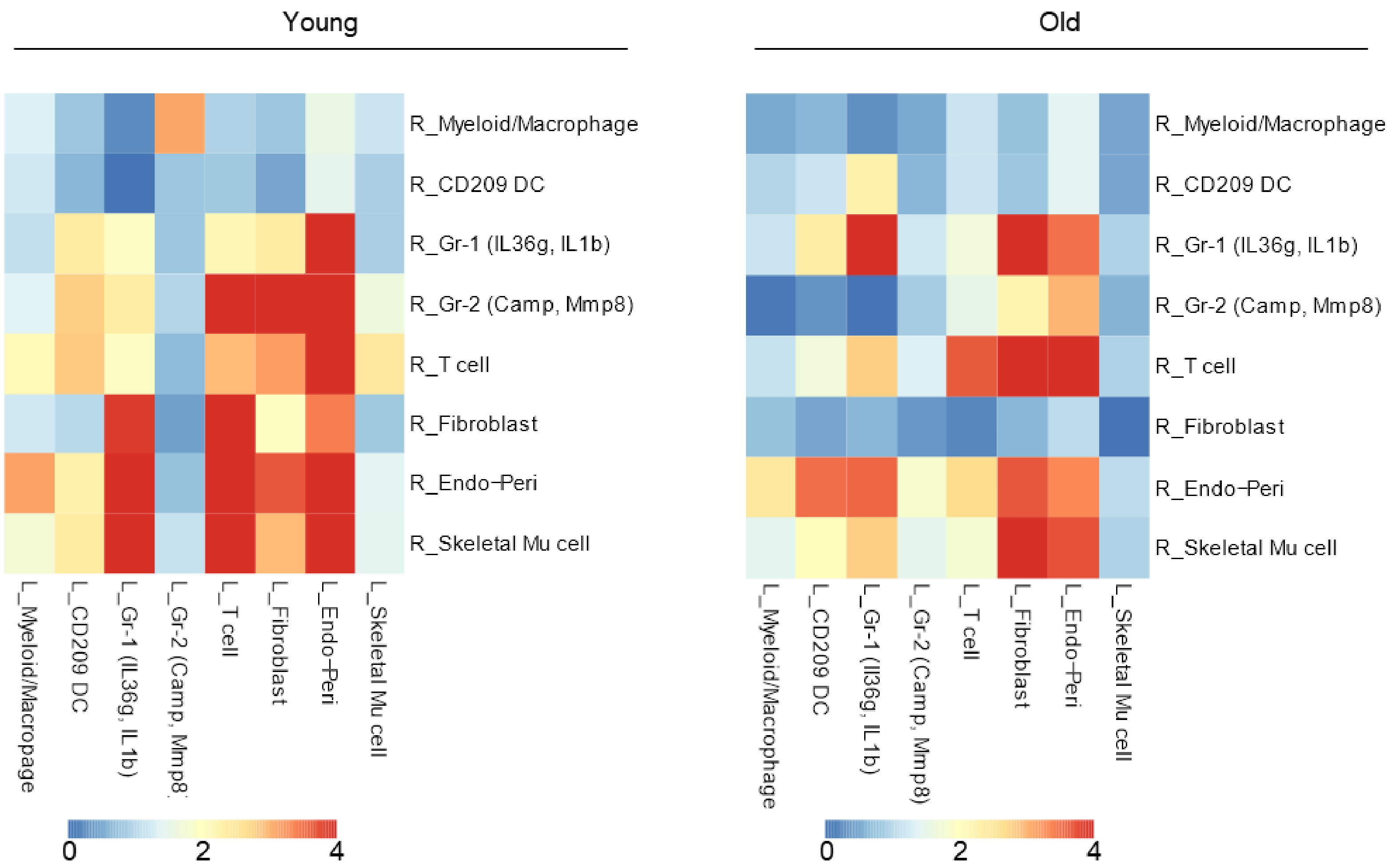


**Supplementary figure 14**

Heatmap of predicted cluster-cluster signaling. The values shown are the summed z-scored expression values for ligands (in the ligand-cluster; L_) targeting receptors predicted to be activated in the receptor-cluster (R_). Higher values indicate increased expression of ligands predicted to be active for a given receptor cluster.

| Generic collagen | Feature score |
|---|---|
| Col3a1 | 9.189499855 |
| Col1a2 | 9.16385746 |
| Sparc | 8.694504738 |
| Malat1 | 7.551837444 |
| Col1a1 | 6.715159416 |
| Fstl1 | 4.818490505 |
| Dcn | 4.776937008 |
| Lgals1 | 3.719181776 |
| Actb | 3.719179153 |
| Tmsb4x | 3.641968966 |
| S100a6 | 3.614605665 |
| Eef1a1 | 3.391069412 |
| Cd63 | 3.276218653 |
| Postn | 3.201486588 |
| Hsp90b1 | 3.167645216 |
| Fn1 | 2.768630981 |
| Hsp90ab1 | 2.576849937 |
| Pabpc1 | 2.459052086 |
| Col5a2 | 2.42209816 |
| Bgn | 2.123658895 |
| Ubb | 2.076804399 |
| Serpinh1 | 1.825502515 |
| Hspa5 | 1.801607609 |
| B2m | 1.752069354 |
| Dpt | 1.749325156 |
| Vim | 1.606626391 |

| Collagen matrix assembly | Feature score |
|---|---|
| Malat1 | 12.1361351 |
| Dcn | 9.362518311 |
| Gsn | 7.329038143 |
| Fos | 6.830287933 |
| Egr1 | 5.194843292 |
| Col1a2 | 5.172890663 |
| Sparc | 4.173226357 |
| Actb | 4.002488613 |
| Jun | 3.907422543 |
| Hspa5 | 3.732988358 |
| Zbtb20 | 3.503760576 |
| S100a6 | 3.088292122 |
| Hsp90b1 | 2.842032909 |
| Ogn | 2.712482214 |
| Klf4 | 2.57933116 |
| Col3a1 | 2.531109333 |
| Fosb | 2.523214817 |
| Hsp90ab1 | 2.477972746 |
| Ubb | 2.413072586 |
| Eef1a1 | 2.337210894 |
| Ftl1 | 2.239730358 |
| Fth1 | 2.198210239 |
| Ptma | 2.172615528 |
| Tmsb4x | 1.959691048 |
| Ahnak | 1.754109383 |
| Serping1 | 1.738965034 |

| Fibroblast progenitor | Feature score |
|---|---|
| Ly6a | 7.413034916 |
| Dpt | 7.394943237 |
| Fstl1 | 6.519348145 |
| Fbn1 | 5.850984573 |
| Fn1 | 5.654873848 |
| Ly6c1 | 5.292335033 |
| Malat1 | 4.941317081 |
| Dcn | 4.908827782 |
| Col3a1 | 4.66126585 |
| C3 | 4.497314453 |
| Plpp3 | 4.289090157 |
| Col14a1 | 4.204661846 |
| Marcks | 3.981361628 |
| Mfap5 | 3.924052715 |
| Col1a2 | 3.898582697 |
| Timp2 | 3.869049072 |
| Sparc | 3.821456909 |
| Sema3c | 3.429731369 |
| Tmsb4x | 3.358593464 |
| S100a6 | 3.232751131 |
| Eef1a1 | 3.14788723 |
| Cd34 | 3.083594799 |
| B2m | 2.967600584 |
| Ogn | 2.885039806 |
| Adamts5 | 2.790465832 |
| Gsn | 2.78393364 |

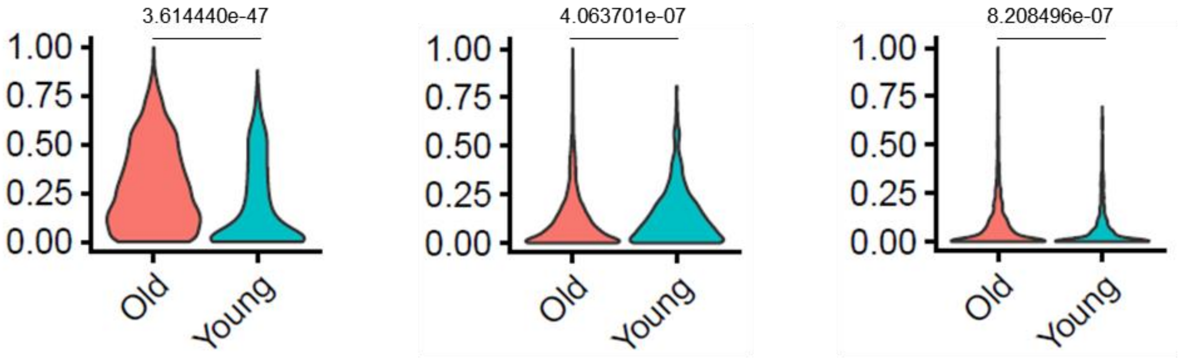


**Supplementary figure 15**

Gene signature of fibroblast populations using NMF CoGAPS. *p* values are determined by Mann-Whitney U test and adjusted with false discovery rate correction for multiple testing.

| Activation cluster | Feature score |
|---|---|
| Ftl1 | 11.4789753 |
| Calm1 | 7.24875927 |
| Crip1 | 7.133933067 |
| Lgals3 | 6.975289822 |
| Gpx1 | 6.546682358 |
| Cyba | 5.309086323 |
| Psap | 5.107288837 |
| Actb | 4.173827648 |
| Ptma | 4.042078972 |
| Arpc1b | 3.912189484 |
| Tpm3 | 3.832166195 |
| S100a6 | 3.582902908 |
| Vim | 3.406517267 |
| Fth1 | 2.901902914 |
| Cst3 | 2.633894682 |
| Anxa2 | 2.517900705 |
| H3f3b | 2.459425211 |
| Cfl1 | 2.385990858 |
| Fcer1g | 2.188590288 |
| Ubb | 2.155832767 |
| Pkm | 2.051351547 |
| Emp3 | 2.041296244 |
| Cstb | 2.026260853 |
| Arpc2 | 2.023267269 |
| Fau | 1.966626763 |
| Tmsb10 | 1.930055499 |

| Regulatory clusters | Feature score |
|---|---|
| Apoe | 9.680174828 |
| Ftl1 | 6.132227421 |
| Fth1 | 5.599677563 |
| C1qb | 5.47872448 |
| H2-D1 | 5.131283283 |
| Itm2b | 5.109345436 |
| Ctsb | 4.965939522 |
| Retnla | 3.533421516 |
| Cxcl2 | 3.477307081 |
| Fcer1g | 3.468292713 |
| C1qa | 3.452937603 |
| Cst3 | 3.392865419 |
| Fos | 3.367647409 |
| Gpx1 | 3.262842894 |
| H2-K1 | 3.202703238 |
| Lgals3 | 3.141122103 |
| Fau | 3.118769407 |
| Ctsd | 2.942102909 |
| C1qc | 2.618610382 |
| H3f3b | 2.587616205 |
| B2m | 2.558019876 |
| Tpt1 | 2.515801907 |
| Ubb | 2.51099658 |
| Pf4 | 2.434630871 |
| Npc2 | 2.393622398 |
| Tyrobp | 2.331031322 |

| Antigen-presentation cluster | Feature score |
|---|---|
| Cd74 | 9.175382614 |
| H2-Aa | 7.442041874 |
| H2-Ab1 | 7.317076206 |
| H2-Eb1 | 5.988512993 |
| Tmsb4x | 3.250351667 |
| Malat1 | 2.900955915 |
| Actb | 2.46359849 |
| Ftl1 | 0.927996814 |
| Cd52 | 0.818355441 |
| Eef1a1 | 0.792192519 |
| B2m | 0.778786957 |
| Lyz2 | 0.730458021 |
| Ctss | 0.688196421 |
| Ptma | 0.672709525 |
| Crip1 | 0.614463389 |
| Calm1 | 0.565919399 |
| Ccr2 | 0.499643713 |
| H2-DMb1 | 0.4949646 |
| Hspa5 | 0.46729821 |
| Cst3 | 0.459704757 |
| Fau | 0.457470298 |
| Pabpc1 | 0.430026203 |
| H2-DMa | 0.422595084 |
| Laptm5 | 0.415646255 |
| Btg2 | 0.39661175 |
| Rack1 | 0.396404803 |

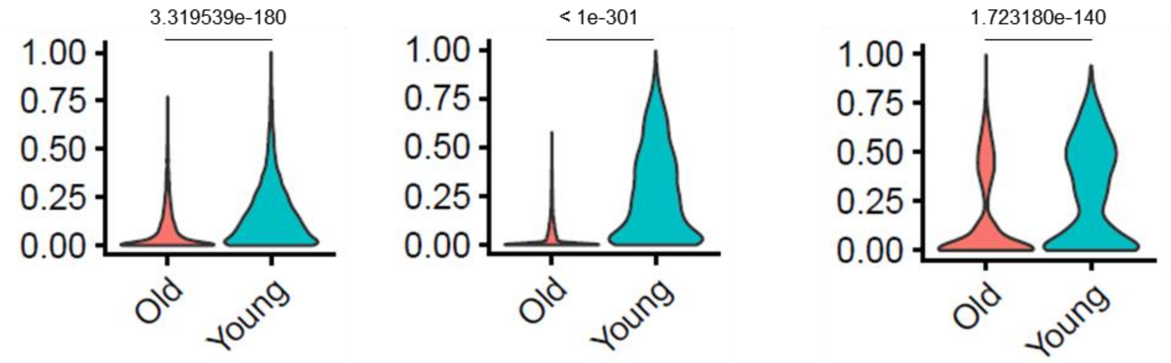


**Supplementary figure 16**

Gene signature of myeloid/macrophage populations using NMF CoGAPS. *p* values are determined by Mann-Whitney U test and adjusted with false discovery rate correction for multiple testing.

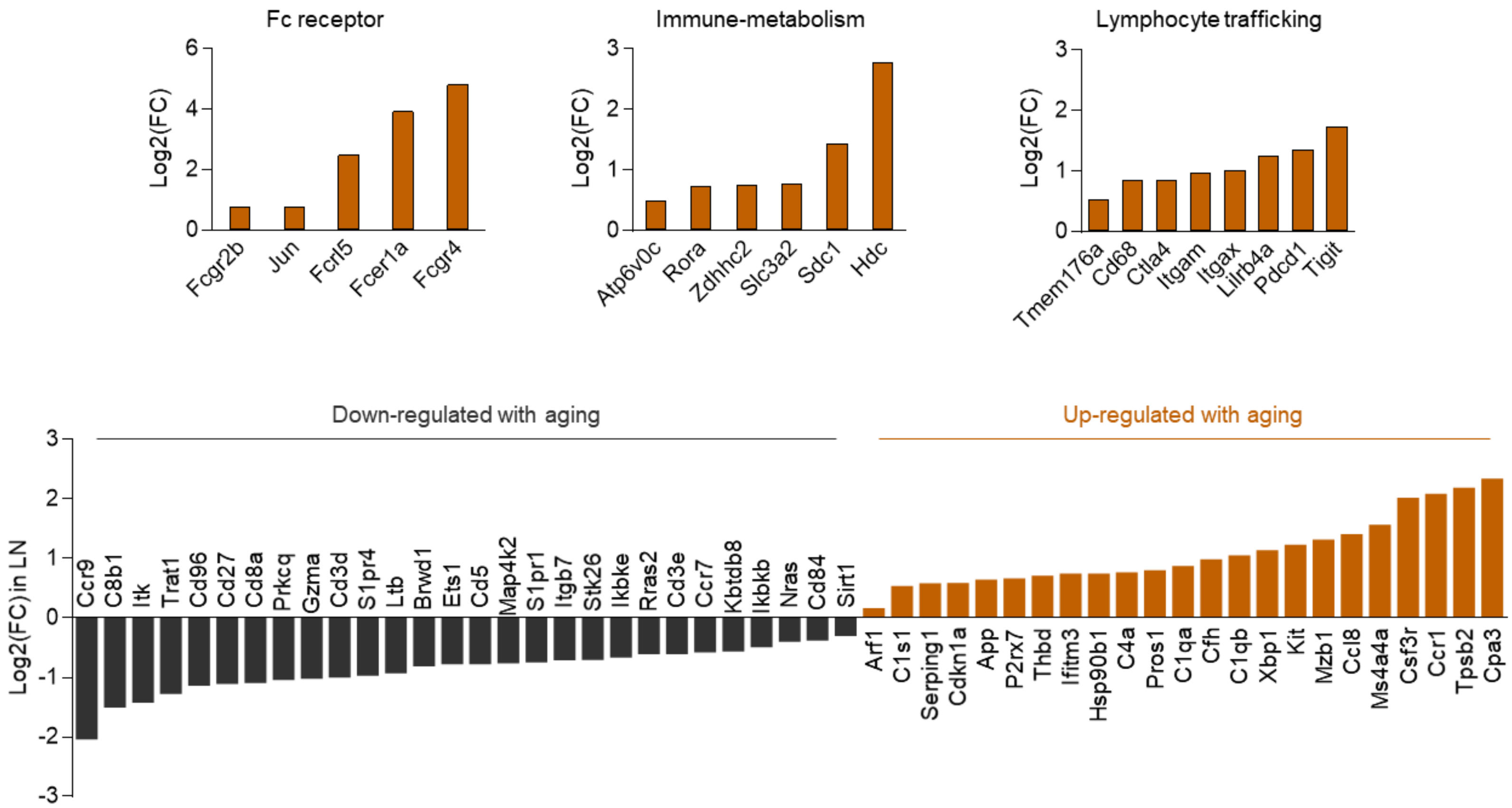


**Supplementary figure 17**

A list of significantly up- or down-regulated genes ($p$<0.05) in the aged lymph nodes compared to young lymph nodes from Nanostring analysis (n=6).

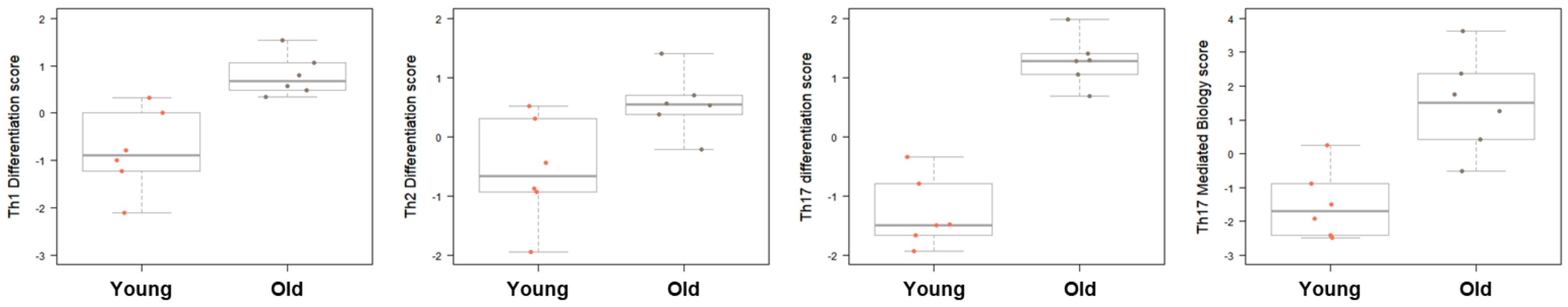


**Supplementary figure 18**

Pathway scoring of the genes associated with Th1-, Th2-, Th17- differentiation or Th17-mediated biology from Nanostring analysis.

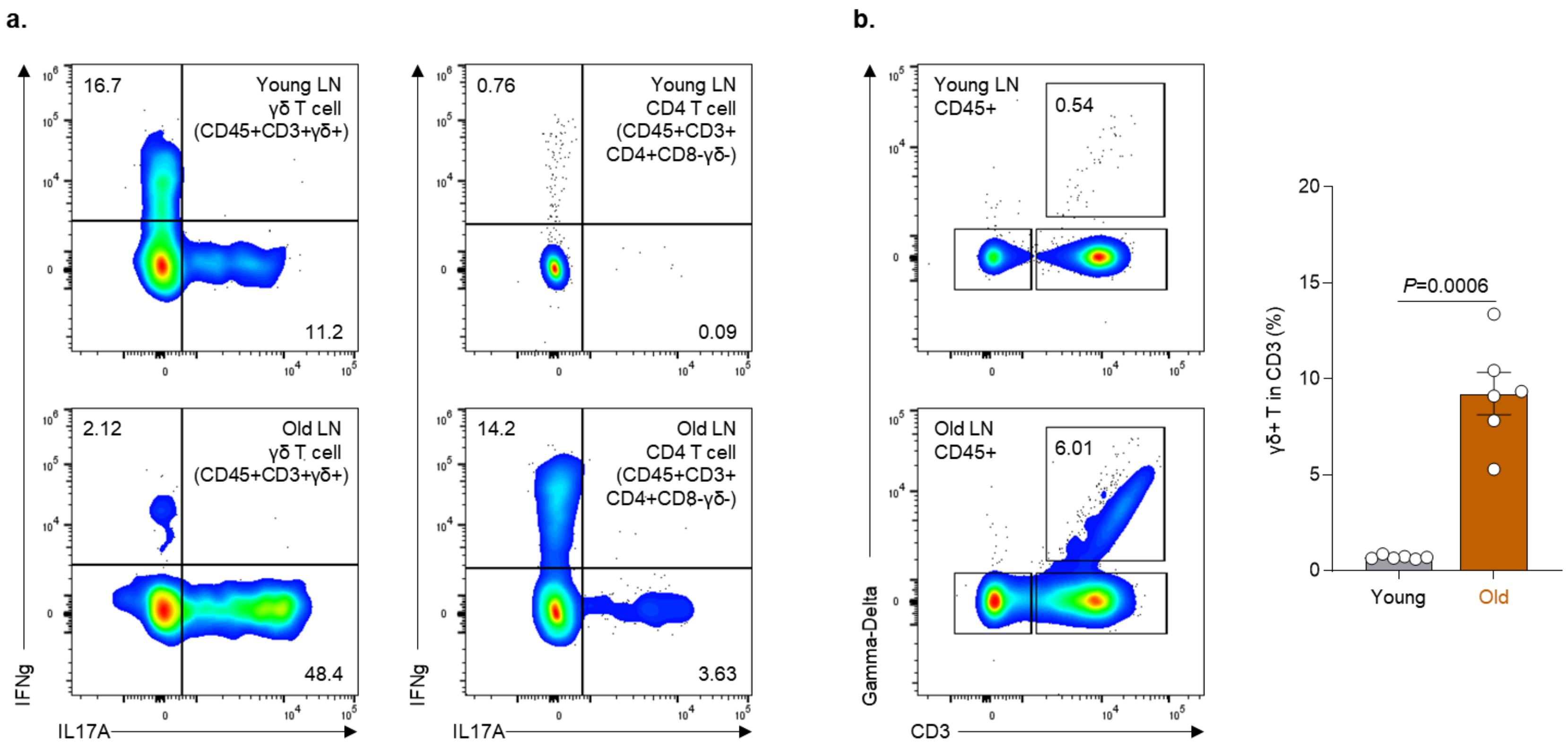


**Supplementary figure 19**

Immune phenotype changes in the inguinal lymph node from young and aged animals without injury or ECM treatment. **a**, Representative images of flow cytometry data showing IFNγ$^{+}$ or IL17A$^{+}$ γδ or CD4 T cells in the lymph node. **b**, Representative images (left), and quantification of flow cytometry data showing γδ T cell population in CD3$^{+}$ T cells in the lymph node (n=6). Unpaired two-tailed t-test (**b)**. For bar graph, data are mean ± s.e.m.

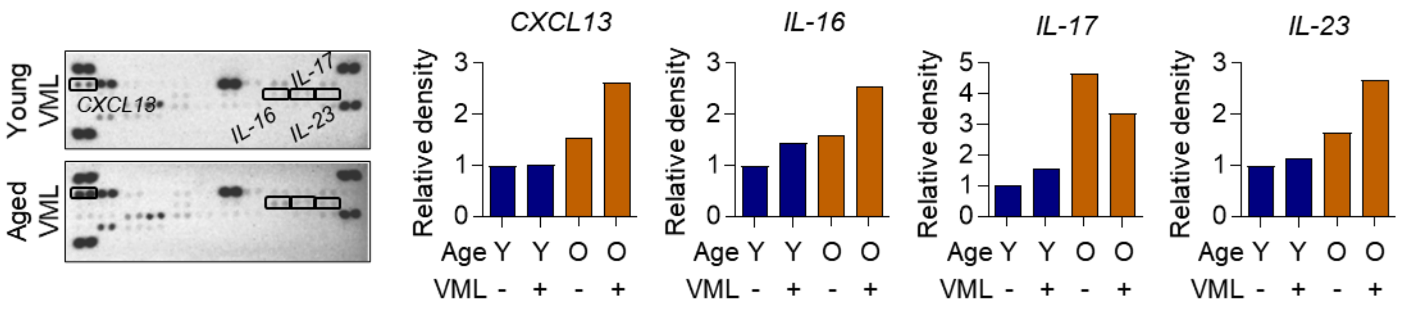


**Supplementary figure 20**

Representative images (left) and quantitative analysis (right) of the proteome profiler array performed on blood serum from young and aged mice 1 week after injury. Protein molecules with significant differences in pixel densities between the groups are labeled and quantified using imageJ. Serum from 3 animals were pooled for analysis.

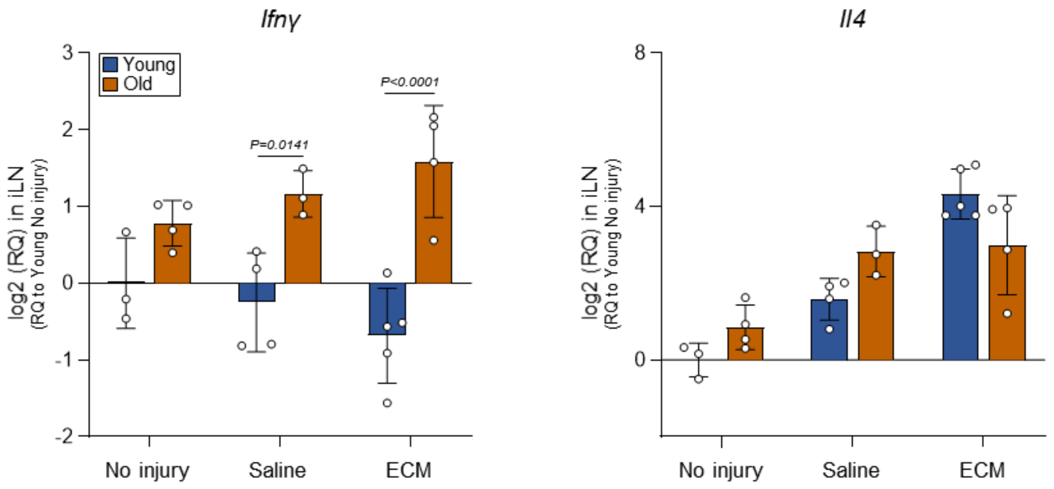


**Supplementary figure 21**

Quantification of Th1 or Th2-related genes in lymph node. Statistical analysis was performed using a two-way ANOVA with Sidak's multiple comparisons test. For all bar graphs, data are mean ± s.d.

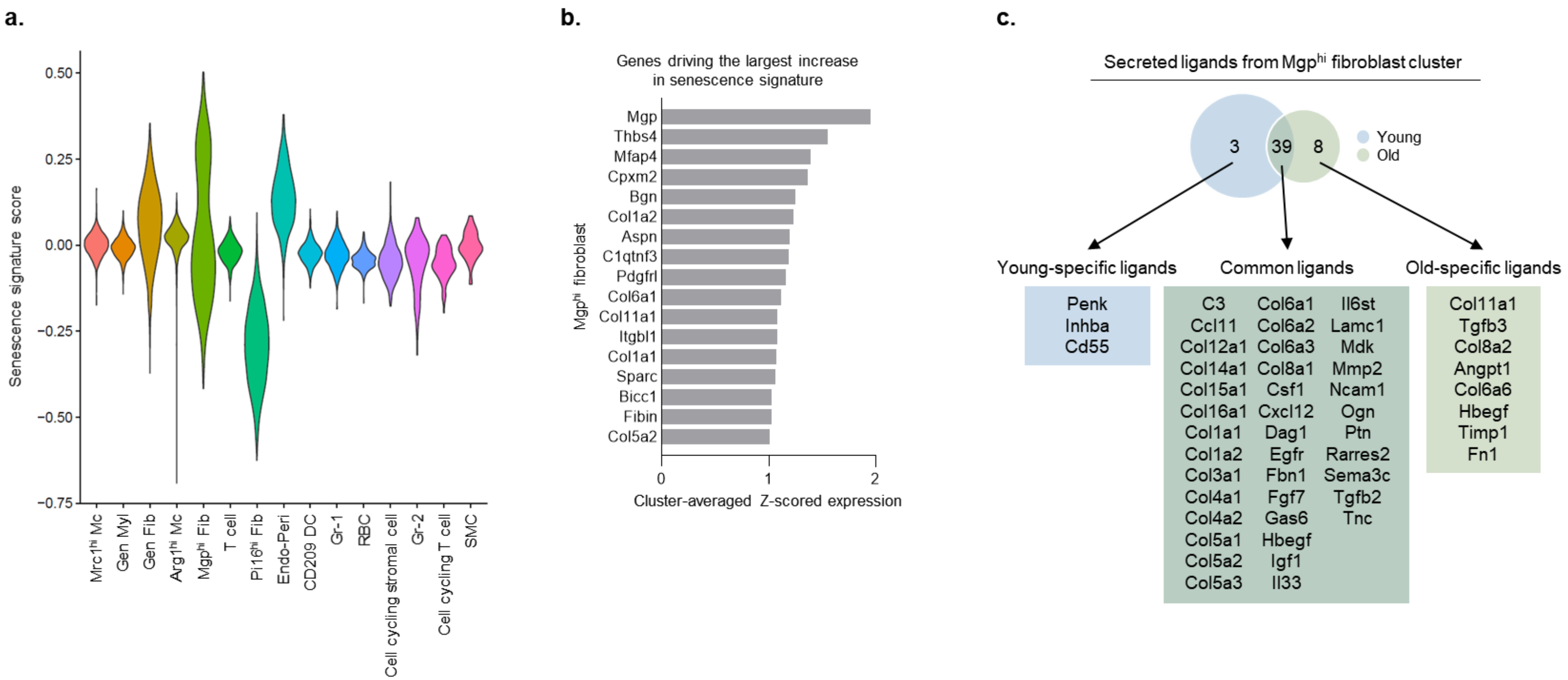


**Supplementary figure 22**

Senescence signature in Mgp$^{hi}$ firoblast cluster. **a,** Senescence scores for cells shown in violin plot grouped by cluster. **b,** Top seventeen gene driving the largest increase in senescence signature in Mgp$^{hi}$ fibroblast cluster. Values are shown as an average z-score expression across cells in the target cluster. **c,** Venn diagram showing a full list of ligands that are common or specific to Mgp$^{hi}$ fibroblast in young and/or old muscle.

**a.**

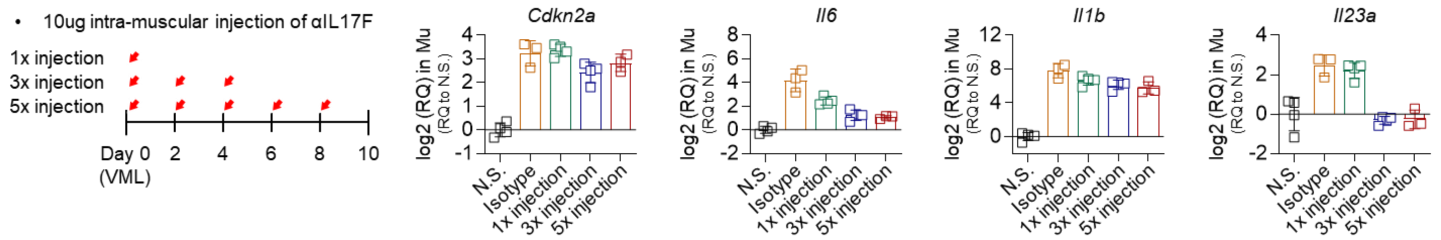


*Cdkn2a*

| Tukey's multiple comparison | | p |
|---|---|---|
| N.S. | Isotype | **** |
| | 1x injection | **** |
| | 3x injection | **** |
| | 5x injection | **** |
| Isotype | 1x injection | NS |
| | 3x injection | NS |
| | 5x injection | NS |
| 1x injection | 3x injection | * |
| | 5x injection | NS |
| 3x injection | 5x injection | NS |

*Il6*

| Tukey's multiple comparison | | p |
|---|---|---|
| N.S. | Isotype | **** |
| | 1x injection | **** |
| | 3x injection | * |
| | 5x injection | NS |
| Isotype | 1x injection | ** |
| | 3x injection | **** |
| | 5x injection | **** |
| 1x injection | 3x injection | * |
| | 5x injection | * |
| 3x injection | 5x injection | NS |

*Il1b*

| Tukey's multiple comparison | | p |
|---|---|---|
| N.S. | Isotype | **** |
| | 1x injection | **** |
| | 3x injection | **** |
| | 5x injection | **** |
| Isotype | 1x injection | NS |
| | 3x injection | * |
| | 5x injection | * |
| 1x injection | 3x injection | NS |
| | 5x injection | NS |
| 3x injection | 5x injection | NS |

*Il23a*

| Tukey's multiple comparison | | p |
|---|---|---|
| N.S. | Isotype | *** |
| | 1x injection | *** |
| | 3x injection | NS |
| | 5x injection | NS |
| Isotype | 1x injection | NS |
| | 3x injection | *** |
| | 5x injection | *** |
| 1x injection | 3x injection | *** |
| | 5x injection | *** |
| 3x injection | 5x injection | NS |

**b.**

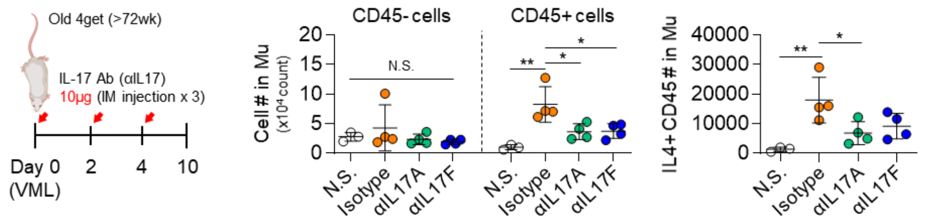


**Supplementary figure 23**

Immune-modulatory effect of αIL17 treatment. **a**, Schematic illustration of experimental design (left) and quantification of senescence- or inflammation-associated genes in muscle after various injection regime (right). **b**, Illustration of experimental design (left) and quantification of flow cytometry data showing the total number of $CD45^-$ or $CD45^+$ live cells or $IL4^+CD45^+$ immune cells in muscle 10 days after VML injury (right). Statistical analysis was performed using a one-way ANOVA with Tukey's multiple comparisons test (n=3-4). *$p$<0.05, and **$p$<0.01. For all bar graphs, data are mean ± s.d.

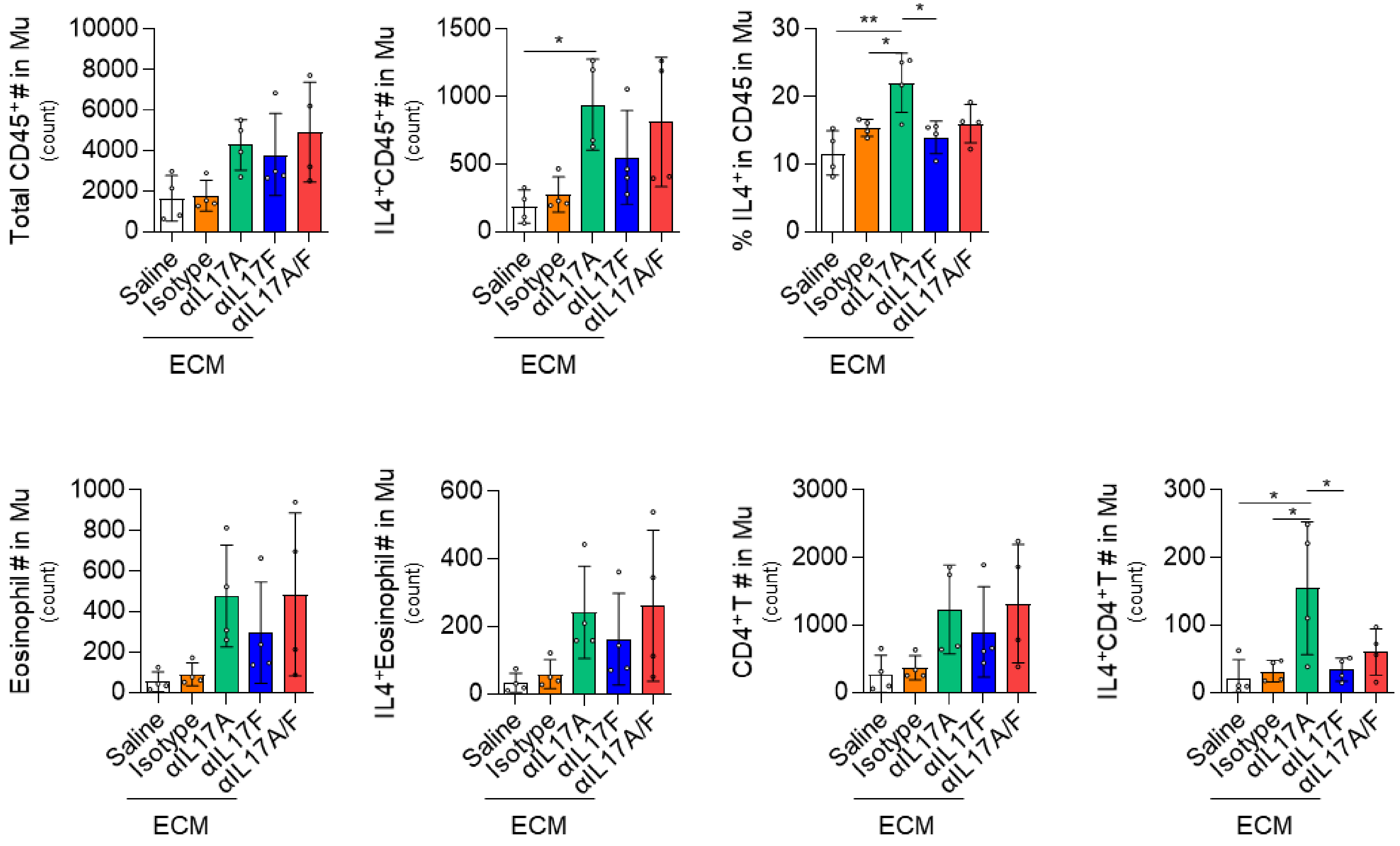


**Supplementary figure 24**

Flow cytometry quantification of the indicated immune cell phenotypes in aged 4Get muscle 6 weeks after injury. Statistical analysis was performed using a one-way ANOVA with Tukey's multiple comparisons test (n=4). *$p<0.05$. For all bar graphs, data are mean ± s.d.

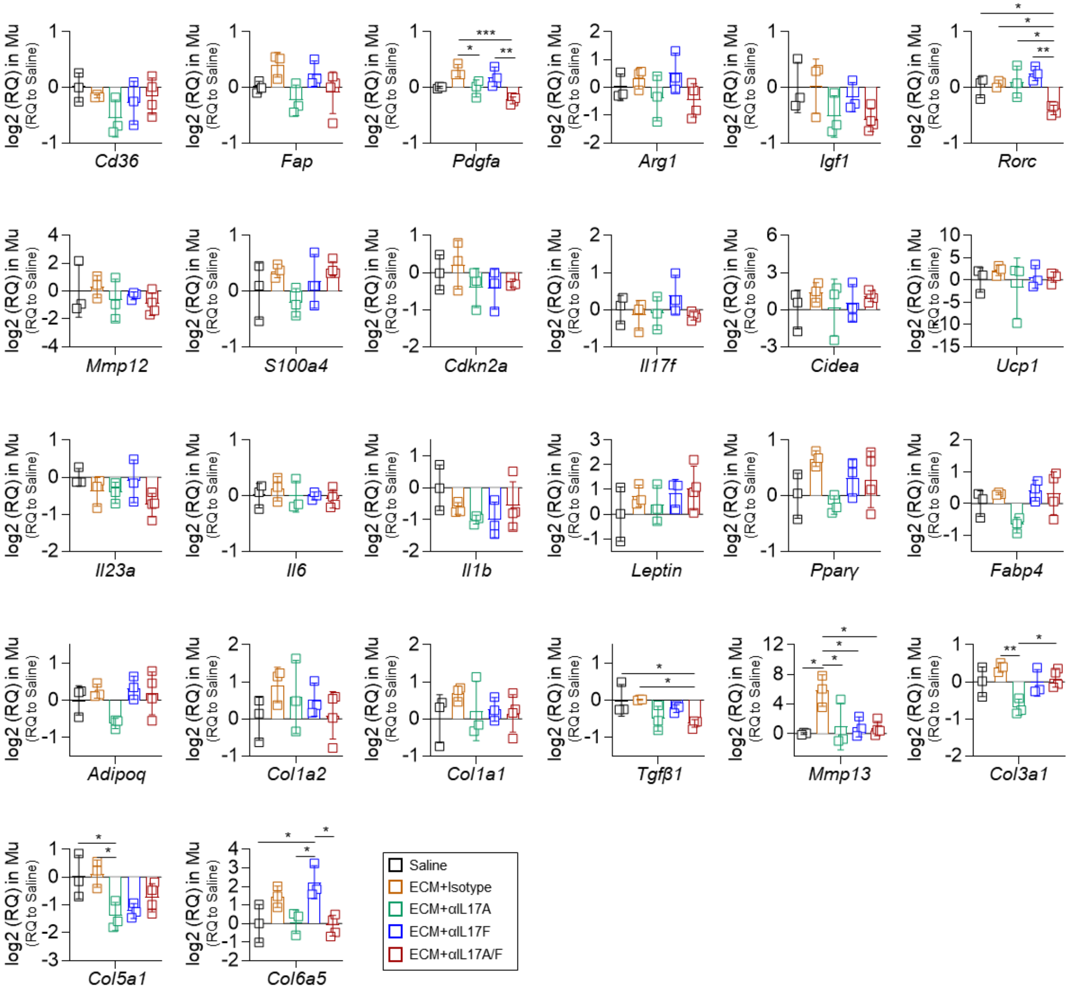


**Supplementary figure 25**

Quantification of inflammation-, senescence-, adipose- or fibrosis-associated genes in aged C57BL/6J muscle 6 weeks after surgery. Statistical analysis was performed using a one-way ANOVA with Tukey's multiple comparisons test (n=3-4). **p*<0.05, ***p*<0.01, ****p*<0.001, and *****p*<0.0001. For all bar graphs, data are mean ± s.d.

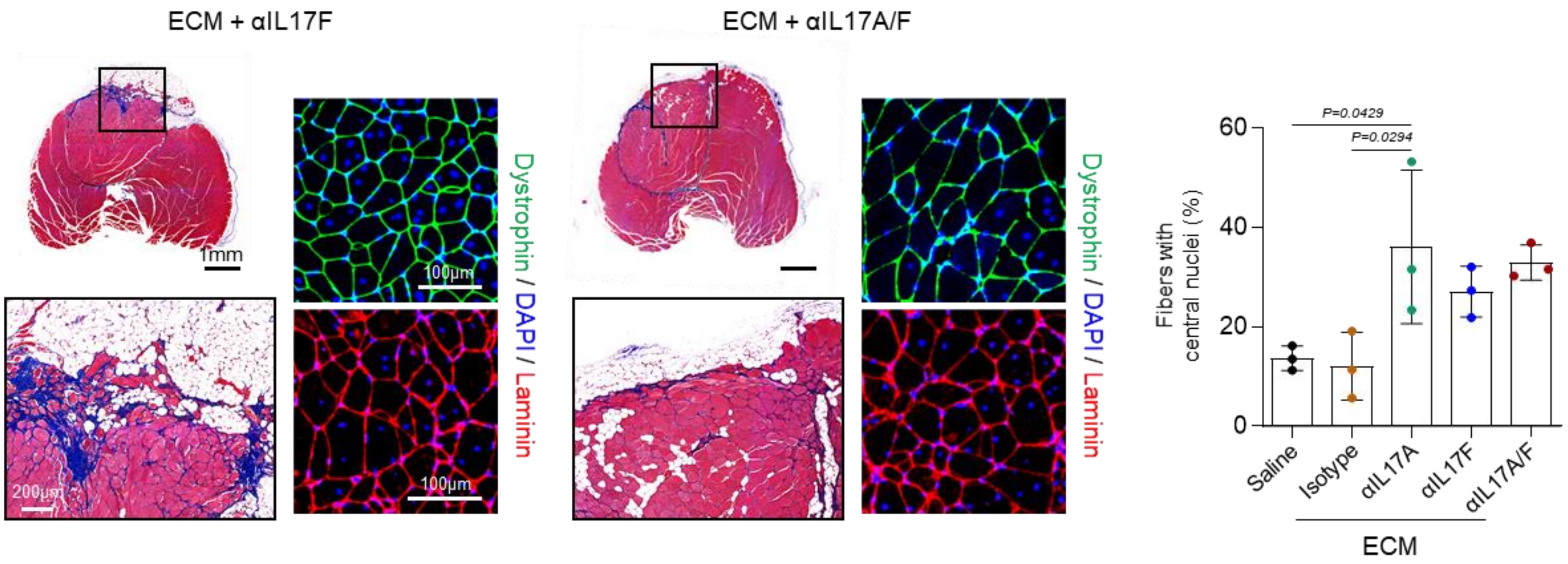


**Supplementary figure 26**

Transverse section of the quadricep muscle 6 weeks after injury stained with Masson's Trichrome, dystrophin or laminin. Quantification of muscle fibers with central nuclei are shown. Statistical analysis was performed using a one-way ANOVA with Tukey's multiple comparisons test. For all bar graphs, data are mean ± s.d.

| Antibodies used for spectral flow cytometry | | |
|---|---|---|
| **Fluorophores** | **Antigen** | **Clone** |
| BV421 | SiglecF | E502440 |
| SuperBright436 | CD19 | 1D3 |
| Pacific Blue | Ly6g | 1A8 |
| BV510 | CD31 | MEC13.3 |
| BV570 | CD45 | 30-F11 |
| BV605 | CD90.2 | 30-H12 |
| BV650 | Ly6c | HK1.4 |
| BV711 | GD | GL3 |
| BV750 | B220 | RA3-6B2 |
| BV785 | F4/80 | BM8 |
| BB515 | cKit | 2B8 |
| SparkBlue550 | CD3 | 17A2 |
| PerCP | MHCII I-A/I-E | M5/114.15.2 |
| BB700 | CD8 | 53-6.7 |
| PerCP-eFluor710 | CD29 | Hmb1-1 |
| PE | CD115 | AFS98 |
| PE-Dazzle594 | CD11c | N418 |
| PE-Cy5.5 | CD68 | C68/684 |
| PE-Cy7 | CD200R3 | BA13 |
| AF647 | NK1.1 | PK136 |
| AF700 | CD11b | M1/70 |
| Zombie NIR | Viability | N/A |
| APC-Fire750 | CD36 | HM36 |
| APC-Fire810 | CD4 | GK1.5 |

| Antibodies used for 4Get mice | | |
|---|---|---|
| **Fluorophores** | **Antigen** | **Clone** |
| GFP | IL4 | N/A |
| PerCP-Cy5.5 | CD3 | 17A2 |
| PE-Cy7 | F4/80 | BM8 |
| PE594 | SiglecF | E502440 |
| PE | CD8 | 53-6.7 |
| APC | CD4 | GK1.5 |
| AF700 | CD11b | M1/70 |
| BV605 | CD45 | 30-F11 |
| BV510 | Ly6c | HK1.4 |
| Pacific Blue | Ly6g | 1A8 |
| eFluor780 | Viability | N/A |

| Antibodies used for IL17A-GFP mice | | |
|---|---|---|
| **Fluorophores** | **Antigen** | **Clone** |
| GFP | IL17A | N/A |
| PE-Cy7 | CD4 | GK1.5 |
| PE594 | GD | GL3 |
| PE | CD3 | 17A2 |
| BV605 | CD45 | 30-F11 |
| Pacific Blue | CD90.2 | 30-H12 |
| eFluor780 | Viability | N/A |

| Antibodies used for intracellular staining | | |
|---|---|---|
| **Fluorophores** | **Antigen** | **Clone** |
| V500 | CD45 | 30-F11 |
| BV711 | CD8 | 53-6.7 |
| AF488 | CD3 | 17A2 |
| PerCP-Cy5.5 | CD19 | 6D5 |
| PE594 | GD | GL3 |
| PE-Cy7 | CD4 | GK1.5 |
| APC | IFNgamma | XMG1.2 |
| AF700 | IL17A | TC11-18H10.1 |
| eFluor780 | Viability | N/A |

| Antibodies used for memory and/or Th17 staining | | |
|---|---|---|
| **Fluorophores** | **Antigen** | **Clone** |
| BV421 | Tbet | 04-46 |
| BV510 | CD45 | 30-F11 |
| BV650 | CD44 | IM7 |
| BV711 | CD4 | GK1.5 |
| SparkBlue550 | CD3 | 17A2 |
| PE | RORγt | Q31-378 |
| PE-Dazzle594 | CD62L | MEL-14 |
| APC-eFluor780 | CD49d | R1-2 |
| Zombie NIR | Viability | N/A |

**Supplementary table 1**

Antibody panel for the flow cytometry analysis.
